# Robust Sequence Determinants of α-Synuclein Toxicity in Yeast Implicate Membrane Binding

**DOI:** 10.1101/2020.05.01.072884

**Authors:** Robert W. Newberry, Taylor Arhar, Jean Costello, George C. Hartoularos, Alison M. Maxwell, Zun Zar Chi Naing, Maureen Pittman, Nishith R. Reddy, Daniel M. C. Schwarz, Douglas R. Wassarman, Taia S. Wu, Daniel Barrero, Christa Caggiano, Adam Catching, Taylor B. Cavazos, Laurel S. Estes, Bryan Faust, Elissa A. Fink, Miriam A. Goldman, Yessica K. Gomez, M. Grace Gordon, Laura M. Gunsalus, Nick Hoppe, Maru Jaime-Garza, Matthew C. Johnson, Matthew G. Jones, Andrew F. Kung, Kyle E. Lopez, Jared Lumpe, Calla Martyn, Elizabeth E. McCarthy, Lakshmi E. Miller-Vedam, Erik J. Navarro, Aji Palar, Jenna Pellegrino, Wren Saylor, Christina A. Stephens, Jack Strickland, Hayarpi Torosyan, Stephanie A. Wankowicz, Daniel R. Wong, Garrett Wong, Sy Redding, Eric D. Chow, William F. DeGrado, Martin Kampmann

## Abstract

Protein conformations are shaped by cellular environments, but how environmental changes alter the conformational landscapes of specific proteins *in vivo* remains largely uncharacterized, in part due to the challenge of probing protein structures in living cells. Here, we use deep mutational scanning to investigate how a toxic conformation of α-synuclein, a dynamic protein linked to Parkinson’s disease, responds to perturbations of cellular proteostasis. In the context of a course for graduate students in the UCSF Integrative Program in Quantitative Biology, we screened a comprehensive library of α-synuclein missense mutants in yeast cells treated with a variety of small molecules that perturb cellular processes linked to α-synuclein biology and pathobiology. We found that the conformation of α-synuclein previously shown to drive yeast toxicity—an extended, membrane-bound helix—is largely unaffected by these chemical perturbations, underscoring the importance of this conformational state as a driver of cellular toxicity. On the other hand, the chemical perturbations have a significant effect on the ability of mutations to suppress α-synuclein toxicity. Moreover, we find that sequence determinants of α-synuclein toxicity are well described by a simple structural model of the membrane-bound helix. This model predicts that α-synuclein penetrates the membrane to constant depth across its length but that membrane affinity decreases toward the C terminus, which is consistent with orthogonal biophysical measurements. Finally, we discuss how parallelized chemical genetics experiments can provide a robust framework for inquiry-based graduate coursework.

## INTRODUCTION

Proteins adopt multiple conformations to facilitate specific biological processes; conversely, protein misfolding can disrupt cellular homeostasis and drive disease. Protein conformation is sensitive to various factors including specific regulatory processes, general homeostatic mechanisms, and chemical/physical properties of the cellular environment. Understanding how cellular factors control protein conformation is critical for determining the basis for normal biological processes and aberrant disease pathologies; however, probing cellular determinants of protein conformation remains challenging due to a dearth of methods that report on protein conformation in living cells.

Deep mutational scanning (DMS) has emerged as a powerful approach for revealing protein conformation in living cells.^1–4^ In DMS, an exhaustive library of protein missense variants is screened for an activity of interest, and the activity of each variant is quantified by deep sequencing.^5^ By determining the pattern of mutations that affects activity, one can build and/or test structural models of proteins. Moreover, this procedure can reveal the conformation(s) that are responsible for an activity of interest.

We recently used DMS to identify biologically active conformation(s) of α-synuclein,^6^ which extensive genetic and pathological evidence implicates as a driver of Parkinson’s disease (PD). Missense mutations^7^ and increases in gene dosage^8^ of α-synuclein are both sufficient to cause inherited, early-onset PD, and the brains of affected individuals accumulate misfolded and mislocalized α-synuclein.^9^ Though predominantly unstructured in solution,^10^ α-synuclein can form an amphiphilic helix,^11^ which allows it to bind to the surface of synaptic vesicles and mediate trafficking dynamics.^12^ At high concentrations, membrane-bound α-synuclein can disrupt lipid membranes, which has been suggested as a mechanism of toxicity in synucleinopathies.^13^ α-Synuclein can also adopt diverse β-sheet-rich aggregates, including amyloids,^14^ which can propagate misfolding in a prion-like manner and drive cell death.^15^ Because multiple structural states of α-synuclein demonstrate toxicity, the conformation(s) that are most responsible for α-synuclein’s toxicity remain a matter of debate.

To address this question, we probed the toxic conformation(s) of α-synuclein in the yeast *S. cerevisiae*,^6^ a validated cellular model of α-synuclein pathobiology.^16^ Though no cellular model can capture the complexities of Parkinson’s disease in humans, which involves a variety of non-cell autonomous processes ultimately leading to systems-level disfunction, expression of α-synuclein in yeast, which lacks synuclein homologs, recapitulates several features of its cellular behavior in neurons, including membrane binding and dose-dependent, gain-of-function aggregation and toxicity.^17^ In addition, the aggregates that develop in yeast consist of clustered, membrane-bound structures,^18^ particularly ER/Golgi vesicles, reminiscent of the composition of Lewy bodies found in PD patients.^19^ This system has also been widely used to identify genetic^20, 21^ and chemical^22^ modifiers of α-synuclein toxicity that have been subsequently validated in neuronal and animal models. α-Synuclein functionally interacts with a variety of fundamental processes of eukaryotic cell biology, including protein production and quality control, endomembrane trafficking, and mitochondria activity, all of which are well conserved between yeast and humans.^16^ Therefore, it is likely that processes perturbed by α-synuclein in yeast will also be affected in human cells.

To identify the conformational state(s) that drive toxicity in yeast, we compared the relative toxicity of 2,600 single missense mutants.^6^ The results of that study inspired our work herein, so we will briefly summarize our previous findings, which led us to propose a membrane-bound helix as the toxic species in yeast (Figure 1). Proline substitutions in the first ~90 residues disrupted toxicity, as did incorporation of polar residues within the membrane-binding face. Toxicity-disrupting mutations recurred with strict 3.67 residue periodicity, consistent with helix formation, and mutational sensitivity decreased toward the C-terminal end of the helix, consistent with increased dynamics in this region.^23, 24^ Therefore, our results indicated that the toxic conformation is a membrane-bound helix with increasing dynamics toward the C terminus. This result is also consistent with previous efforts to identify missense suppressors of α-synuclein toxicity in yeast, which likewise identified membrane-binding capacity as a critical determinant of toxicity.^25^ Interestingly, we did not observe any features that would suggest aggregation or β-sheet structure in the toxic conformation.

**Figure 1.**
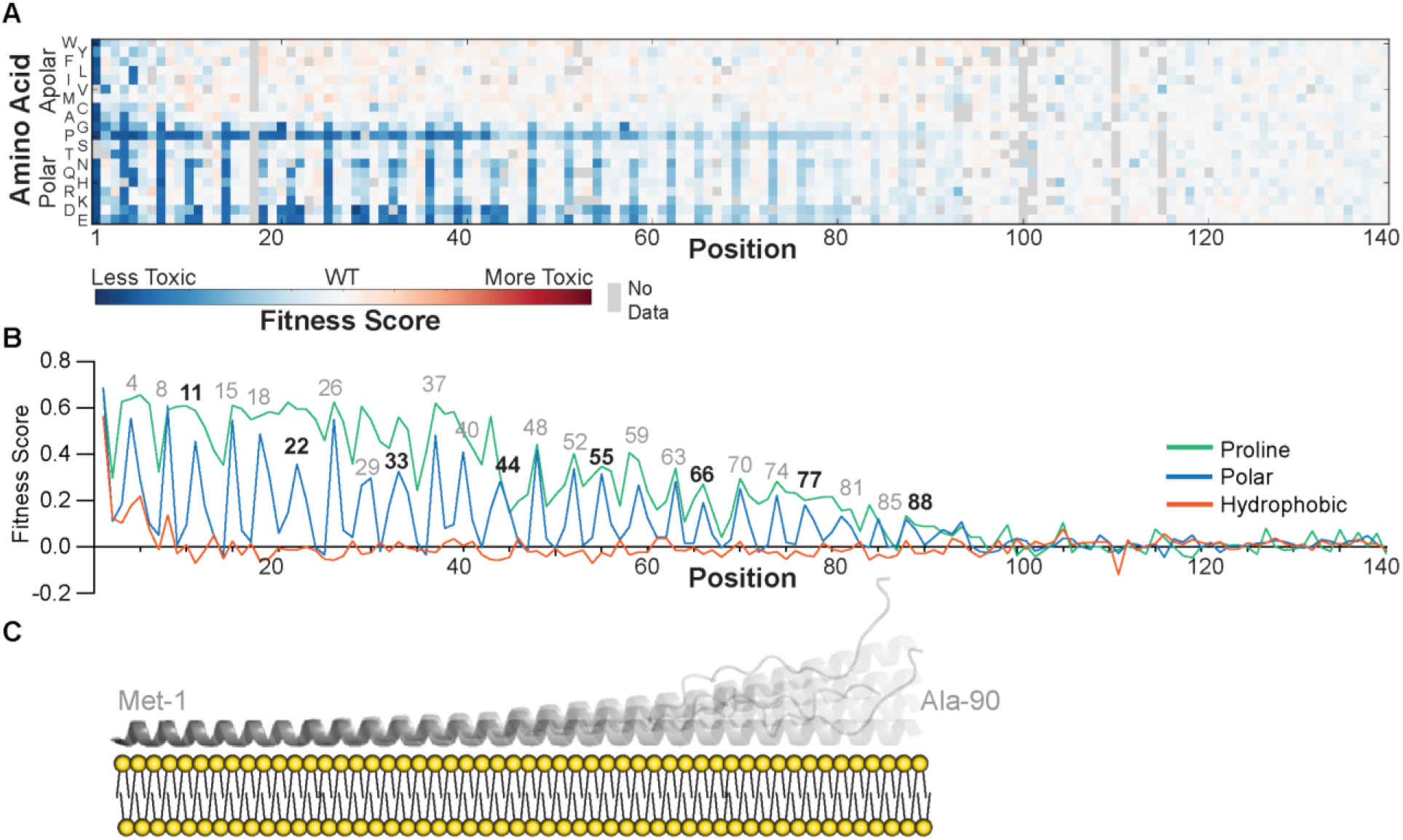
Previously mapped sequence–toxicity landscape of α-synuclein identifies a toxic molecular conformation.^6^ (A) Fitness scores (defined as the slope of the line describing change in log-transformed variant frequencies over time, represented by a red-white-blue color scale) for expression of α-synuclein missense mutants in yeast.^6^ (B) Average fitness scores of mutants with hydrophobic (W, Y, F, L, I, V, M, C, A; red), polar (S, T, N, Q, H, R, K, D, E; blue), or proline (green) residues.^6^ (C) The structural model derived from the sequence–toxicity landscape in panel A: an extended, membrane-bound 11/3-helix with increasing dynamics toward the C terminus.^6^ These results were obtained previously^6^ and are revisited here to provide context for the effects of cellular perturbations on the fitness landscape.

Because α-synuclein adopts a variety of structures, we were intrigued that the sequence landscape was consistent with a single toxic conformation; however, it is possible that additional species contribute to toxicity downstream of the membrane-bound helix. Moreover, several lines of evidence suggest that the cellular environment might alter α-synuclein’s conformational landscape,^26^ including the sensitivity of intrinsically disordered proteins to environmental conditions,^27^ the effect of environmental stresses on α-synuclein toxicity,^28^ the cell-type selective vulnerability in PD,^29^ the age-dependence of α-synuclein pathology,^30^ and the variety of α-synuclein species that demonstrate toxicity in other models.^15, 31, 32^ Given the complex conformational landscape of α-synuclein and known environmental contributions to toxicity, we hypothesized that cellular stresses might create a less permissive folding environment that would either perturb the structure of the toxic species or reveal contributions from species whose toxicity is not evident under unstressed conditions. For example, interventions by the proteostasis network might resolve otherwise disruptive species (e.g., oligomers or amyloids) or directly remodel the active conformation. Alternatively, the cell might harbor multiple sensitivities to α-synuclein, some of which might dominate in different environmental conditions. This would be consistent with the likely scenario that α-synuclein exerts toxicity through multiple mechanisms; because only a small percentage of synucleinopathy cases have clear genetic etiologies, environmental stresses likely predispose cells to particular mechanisms of toxicity.

If additional conformational states contribute to toxicity under different environmental conditions, a variety of changes to the sequence–toxicity landscape might be observed. For example, the formation of β-sheet structure would cause a change in the periodicity of the disrupting mutations. Increased toxicity due to oligomers or amyloids should result in increased correlation of toxicity data with data from aggregation or amyloid prediction algorithms. Formation of specific protein–protein interfaces should manifest in positions sensitive to the steric bulk of substitution. In addition, if multiple conformations were to contribute to toxicity simultaneously, a superposition of sequence landscapes might result, which could be resolved by subtracting out mutational signatures associated with known conformational states. Mutational signatures for separate, coexisting toxic conformations might also be resolved based on their relative intensities. For example, if fitness scores fall into discrete high-, intermediate-, and low-fitness subpopulations, the features of the sequence landscape for each subpopulation might reveal distinct conformational signatures. In theory, the detection of alternative conformations is limited by the noise in the sequence–toxicity landscape of the dominant conformation; in this regard, the relatively high reproducibility between experimental replicates (Figure S13) underscores the sensitivity of our approach. Nonetheless, some conformations, such as those that are robust to single amino-acid substitutions, are unlikely to be revealed by deep mutational scanning and would not be evident in the fitness landscape of perturbed cells.

To address these hypotheses, we used small molecules to alter cellular homeostasis and create distinct folding environments; we then used DMS to read out quantitative changes in the sequence determinants of the toxic conformational state of α-synuclein. DMS is an ideal approach for these questions because structural information can be inferred from phenotypic measurements obtained in living cells. The parallel nature of these experiments also offers an opportunity for authentic inquiry in a classroom-based setting; these studies provide training in critical thinking and creativity within a supportive environment that builds leadership and cross-disciplinary engagement.^33^ We have previously undertaken similar course-based experiments examining environmental constraints on ubiquitin function;^34, 35^ therefore, to sustain the educational impacts of this initiative and ensure authentic, open-ended inquiry, we undertook our experiments on α-synuclein in a project-based course for first-year biophysics and bioinformatics graduate students in the UCSF Integrative Program for Quantitative Biology. The resulting course thereby provided a unique training opportunity integrating experimental and computational approaches to enable both hypothesis-driven and unbiased inquiry.

## RESULTS AND DISCUSSION

### Chemical genetics approach to revealing additional active states

To probe additional conformational states that contribute to α-synuclein toxicity, we perturbed a variety of cellular processes using small molecules and identified ensuing changes in the sequence–toxicity landscape using DMS (Figure 2). We selected cellular stresses relevant to α-synuclein pathobiology (Table 1), including perturbations to vesicle trafficking,^20, 21^ oxidative stress,^28^ the unfolded protein response,^36^ protein degradation and autophagy,^37, 38^ chaperone activity,^39^ and membrane composition.^40^

**Figure 2.**
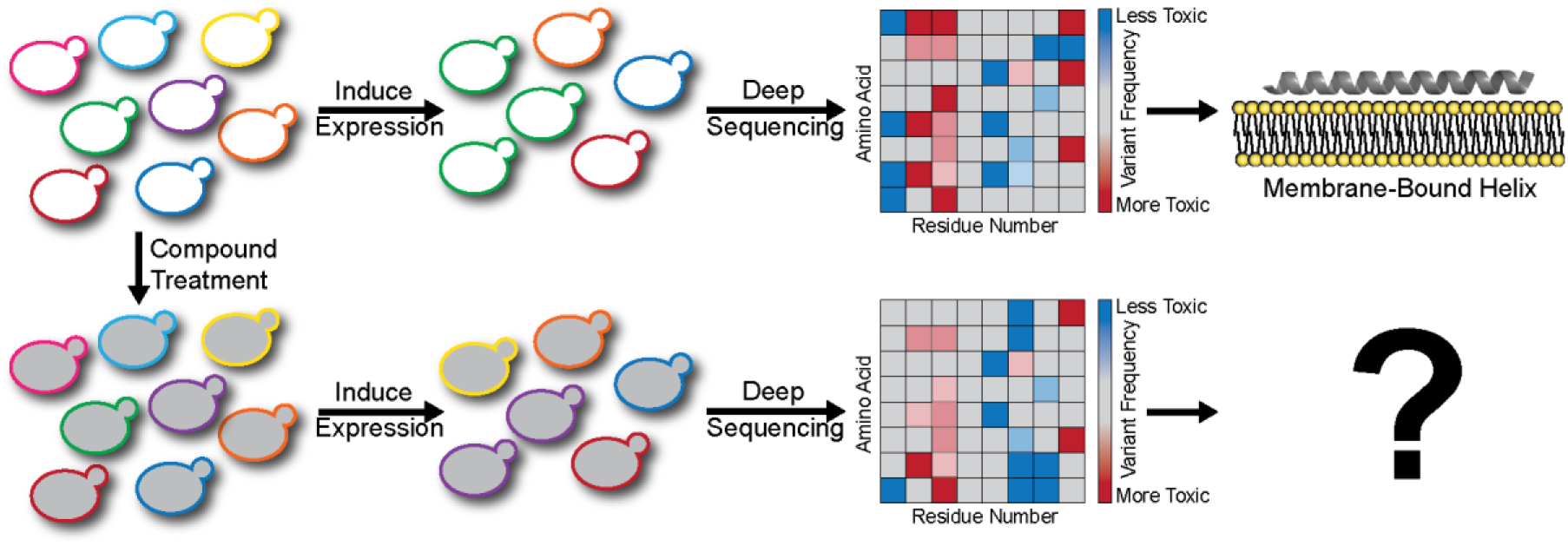
Chemical genetics approach to probe sequence and structural determinants of α-synuclein toxicity. A pooled library of yeast cells, each expressing a different missense variant of α-synuclein (represented by yeast cells with differently colored outlines), is treated with a variety of small molecules to perturb cellular proteostasis (represented by different intracellular shading [i.e., white vs. grey]). Induction of α-synuclein expression creates a selective pressure that causes changes in the relative frequency of each missense variant in the population. This change in frequency can be quantified by counting the number of occurrences of each variant in the population over time using deep sequencing. The resulting sequence–toxicity landscape, where the relative toxicity of each variant is represented by a red-white-blue color scale, reveals structural features that can implicate specific conformational states as drivers of toxicity.

**Table 1.** Effects of chemical perturbations

| **Table 1.** Effects of chemical perturbations |  |  |  |  |
| --- | --- | --- | --- | --- |
| **Compound** | **Target and Mode of Action** | **Primary Cellular Effects** | **Connection to α-Synuclein** | **Hypothesized Effects on α-Synuclein Fitness Landscape** |
| Dopamine | Covalent modification of α-synuclein<sup>41</sup> | Neurotransmitter | α-Synuclein deficiency alters dopamine signaling;<sup>12</sup> dopaminergic neurons degenerate selectively in PD;<sup>29</sup> dopamine promotes oligomer formation, inhibits fibrillization<sup>41</sup> | Residues that interact with dopamine will show disruptive mutations selectively in the presence of dopamine; sequence signatures of oligomers (e.g., positions sensitive to steric hindrance) will emerge |
| Menadione | Quinone reduction by molecular oxygen generates superoxide radicals | Induces oxidative stress | Various α-synuclein species induce oxidative stress;<sup>42</sup> oxidative modification of α-synuclein is observed in patients<sup>43</sup> and inhibits fibrillization;<sup>44</sup> menadione exacerbates α-synuclein toxicity in yeast<sup>45</sup> | Residues whose oxidation contributes to α-synuclein toxicity will become more sensitive to mutation; compromised capacity to resolve oxidative stress might reveal conformations that drive toxicity by increasing oxidative stress |
| Melatonin | Scavenges free radicals | Relieves oxidative stress | Various α-synuclein species induce oxidative stress;<sup>42</sup> oxidative modification of α-synuclein is observed in patients<sup>43</sup> and inhibits fibrillization;<sup>44</sup> melatonin attenuates α-synuclein toxicity in yeast challenged by menadione<sup>45</sup> | Residues that become modified by oxidative stress will be less sensitive to mutation; enhanced resilience to oxidative stress might reveal conformations that drive toxicity through other mechanisms |
| Miconazole | Inhibits 14α-sterol demethylase, reducing ergosterol production<sup>46</sup> | Decreases membrane sterol content, affecting membrane fluidity and integrity<sup>46</sup> | Sterol content modulates the membrane-binding affinity of α-synuclein<sup>47</sup> | Membrane-contacting residues will be selectively affected; cells might maintain resistance only to variants with the weakest membrane affinity |
| Brefeldin A | Inhibits COP-I coat assembly | Disrupts Golgi-ER and other membrane transport processes; induces the unfolded protein response | ER-Golgi trafficking genes modify α-synuclein toxicity;<sup>20</sup> α-synuclein inclusions involve clustered ER/Golgi vesicles<sup>18, 19</sup> | Because cells will be more sensitive to agents that interfere with trafficking, cells should be more sensitive to all variants except those with the weakest membrane affinity, which are less likely to affect trafficking; sequence signatures of alternatively localized α-synuclein should emerge |

| **Table 1.** Effects of chemical perturbations (continued) |  |  |  |  |
| --- | --- | --- | --- | --- |
| **Compound** | **Target and Mode of Action** | **Primary Cellular Effects** | **Connection to α-Synuclein** | **Hypothesized Effects on α-Synuclein Fitness Landscape** |
| MG-132 | Covalent inhibitor of the 20S proteasome | Reduces degradation of ubiquitin-modified proteins | Lewy bodies are enriched in ubiquitinated proteins, including α-synuclein;<sup>48</sup> ubiquitination is a common cellular strategy for mitigating misfolded proteins; α-synuclein impairs proteasome function<sup>49</sup> | Residues whose ubiquitination modifies α-synuclein toxicity will be less sensitive to mutation; species that are more efficiently ubiquitinated and cleared will become apparent |
| Tunicamycin | Inhibits glycosyl-transferases involved in glycoprotein synthesis<sup>50</sup> | Creates a burden of misfolded ER proteins; induces the unfolded protein response<sup>50</sup> | α-Synuclein accumulation induces the unfolded protein response;<sup>21</sup> α-synuclein interferes with glycoprotein maturation;<sup>21</sup> tunicamycin induces α-synuclein oligomerization in cellular models<sup>36</sup> | Cells will become less resistant to all variants except those with the weakest membrane affinity, which are less likely to affect glycoprotein maturation; additional sequence determinants of oligomerization might appear; sequence determinants of species resolved by the unfolded protein response will appear |
| Geldanamycin | ADP/ATP-competitive inhibitor of Hsp90<sup>51</sup> | Reduces proteostatic capacity<sup>51</sup> | Hsp90 reduces α-synuclein aggregation and toxicity by modifying oligomers<sup>52</sup> | Sequence signatures of conformations resolved by Hsp90 will appear; residues that directly interact with Hsp90 will be less sensitive to mutation |
| Rapamycin | Inhibitor of mTORC1 formation<sup>53</sup> | Inhibits translation; induces autophagy<sup>53</sup> | α-Synuclein aggregates are cleared by autophagy;<sup>37</sup> α-synuclein aggregation impairs autophagy<sup>54</sup> | Variants with sub-saturating toxicity will increase in fitness due to reduced translation; residues that mediate trafficking to autophagosomes will be more sensitive to mutation; sequence signatures of toxic species not resolved by autophagy will be revealed |
| Spermidine | Inhibits histone acetyltransferases, altering the epigenetic state of histone H3 and inducing transcription of autophagy-related genes<sup>55</sup> | Induces autophagy; suppresses oxidative stress<sup>55</sup> | α-Synuclein aggregates are cleared by autophagy;<sup>37</sup> synuclein aggregation impairs autophagy<sup>54</sup> | Residues that mediate trafficking to autophagosomes will be more sensitive to mutation; sequence signatures of toxic species not resolved by autophagy will be revealed |

We treated our yeast library with compound concentrations that reduced growth rate, where possible, to confirm that the compound was active in cells. Compounds that failed to induce a growth defect (dopamine, melatonin, and spermidine) were employed at high concentrations previously reported to perturb survival processes in yeast.^45, 55, 56^ Each population was grown for several generations, and the change in relative frequency of each variant in the population over time was monitored by deep sequencing. We then compared the resulting sequence–toxicity landscape to that obtained in untreated controls.

### Retention of structure upon diverse chemical treatments

To determine the effect of chemical perturbations, we compared fitness scores for each mutant in the presence versus the absence of a given compound. If the compound had no effect on the conformation responsible for toxicity, the data would retain periodicity reflecting the α-helix, and the introduction of helix-breaking residues such as proline would suppress toxicity throughout the helical region. This is precisely what is observed (Figures 3, S1–S10).

**Figure 3.**
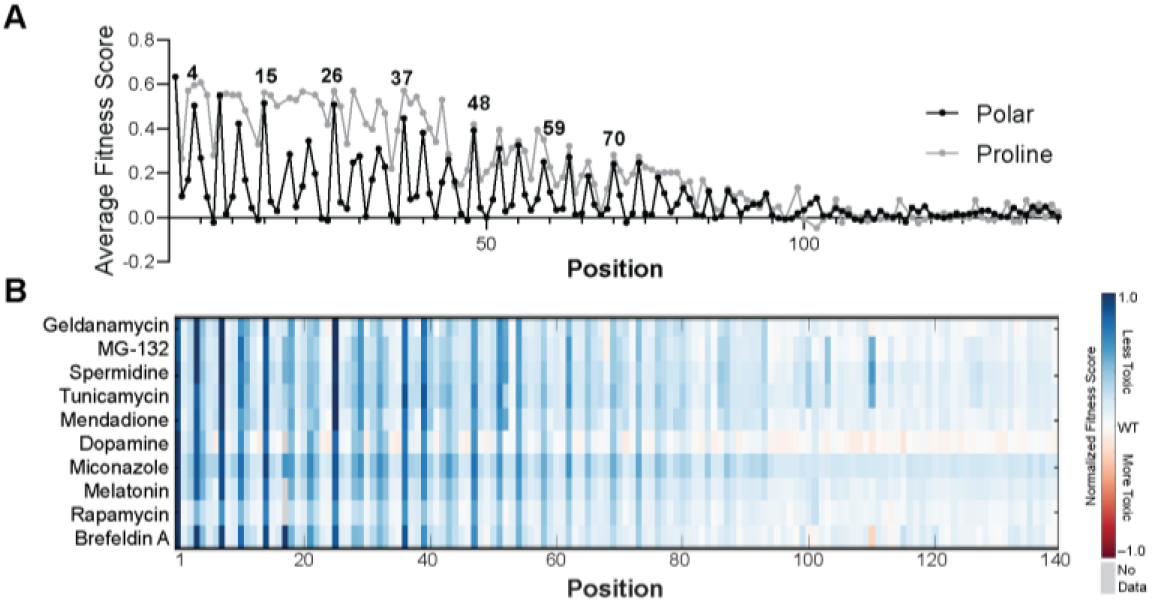
Conformational signatures of toxic α-synuclein species in yeast are robust to diverse chemical perturbations. (A) Average fitness scores of mutants with polar (S, T, N, Q, H, R, K, D, E) or proline residues in untreated cells.^6^ These data were collected previously and are shown here for comparison. (B) Average fitness scores of mutants with polar residues when assayed in yeast cells perturbed by small molecules. Each column reflects the average fitness scores of mutants with polar substitutions at a given position. Each row reflects the fitness scores for a different treatment. Fitness scores are normalized for comparison and shown as a red-white-blue color scale. Across all conditions, α-helical periodicity in the mutational signature is retained.

In each condition, polar residues continue to increase fitness when substituted on the apolar face of α-synuclein’s membrane-binding helix (Figure 3). Moreover, the effects of apolar-to-polar and proline substitutions decrease along the chain in each condition. We therefore conclude that the same dynamic membrane-binding helix is the key driver of toxicity over a wide range of chemical perturbations.

### Chemical perturbation modulates efficiency of single-site suppressors of toxicity

To determine the impact of chemical perturbation on the efficiency with which missense mutations suppress α-synuclein toxicity, we compared the fitness scores measured in the presence and absence of each compound. If treatment places a stress on the system that globally increases sensitivity to α-synuclein, then it should become more difficult to suppress toxicity with otherwise beneficial substitutions (e.g., apolar-to-polar substitutions), resulting in a strong correlation between control and treatment, but with a slope less than one. Conversely, if the chemical perturbation induces processes (e.g., induction of beneficial chaperones or proteases) that rescue the toxic effects of α-synuclein, then a slope greater than one should be observed. Finally, outliers that deviate significantly from the linear trend would reveal any sequence-specific effects on toxicity caused by the treatment.

The data for DMS in the presence of menadione, melatonin, spermidine, dopamine, miconazole, and MG-132 were nearly perfectly correlated with the control and had slopes near one (Figure 4). Thus, these conditions had no global effect on the sensitivity of cells to the expression of α-synuclein. By contrast, the correlations between data for controls and data for tunicamycin, geldanamycin, brefeldin A, and rapamycin had significantly reduced slopes. The relative rates of change in frequency for mutants vs. wild-type (WT), both in the presence of a given compound, were compared to account for the fact that compound toxicity limits growth rate under these conditions (Figure S12). A second effect seen in plots with the slopes significantly less than one was the emergence of a large cluster of mutants with a fitness score near zero for the drug-treated population (e.g., brefeldin A), indicating that these mutants had significant effects under normal conditions but were unable to impact fitness in the presence of this chemical.

**Figure 4.**
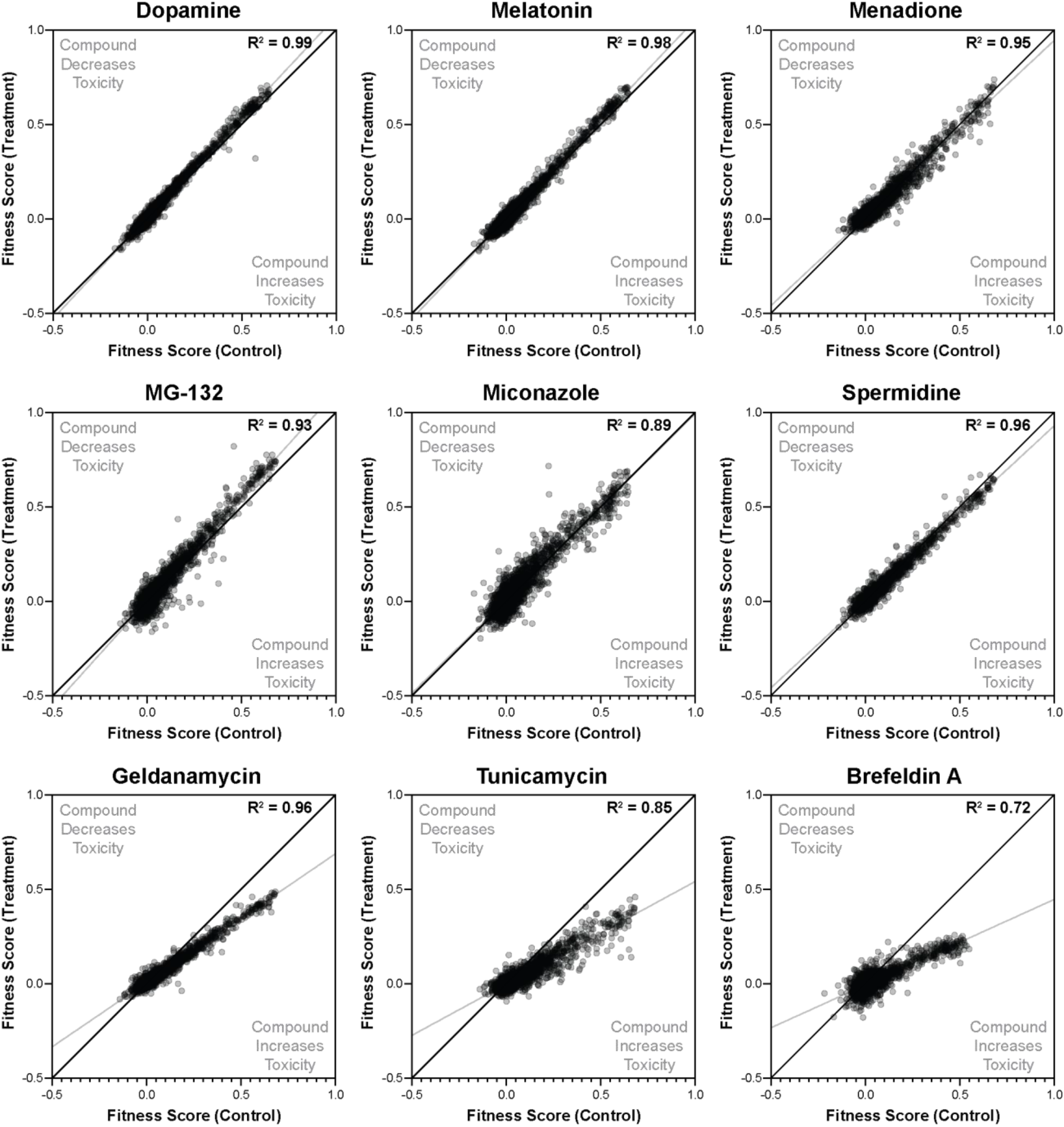
Fitness scores of yeast expressing α-synuclein missense variants obtained in the presence of the compounds in Table 1, relative to those obtained in untreated controls.

The fitness scores we obtained in the presence of this panel of treatments were remarkably robust. We probed for systematic changes in the fitness scores of various subsets of mutants, including mutants with high/low toxicity, variants with substitutions on particular faces of the amphiphilic helix, variants with substitutions at any specific position, or mutants substituted with a particular amino acid; however, we identified no significant changes in the fitness scores of any of these subsets. We also examined the identities of variants with “outlying” fitness scores, but these values had high uncertainties (due primarily to low library coverage) and there were no clear features shared between them. In addition, we probed for changes in periodicity of the mutations that suppress toxicity and correlation between toxicity data and expectations from aggregation/amyloid prediction algorithms, but again, we were unable to establish statistical support.

Among the ten chemical perturbations we tested, we identified one compound, rapamycin, that displayed non-linear behavior between treated and untreated conditions, suggesting the presence of mutant-specific effects of rapamycin treatment. Specifically, although the entire set of mutants are poor suppressors of toxicity in the presence of rapamycin, two subsets of variants showed a small but significant positive deviation from the linear relationship between fitness scores observed in the presence versus absence of rapamycin: (1) those with substitutions of Met-1 (Figure 5A), which dramatically reduce expression of α-synuclein,^6^ and (2) those with acidic residues on the hydrophobic face of the amphiphilic helix (Figure 5B/C). We showed previously that incorporation of polar residues on the hydrophobic face dramatically reduces membrane binding,^6^ so one feature shared amongst the variants whose fitness was modified by rapamycin is decreased membrane-bound α-synuclein, either by reduced expression or reduced membrane affinity.

**Figure 5.**
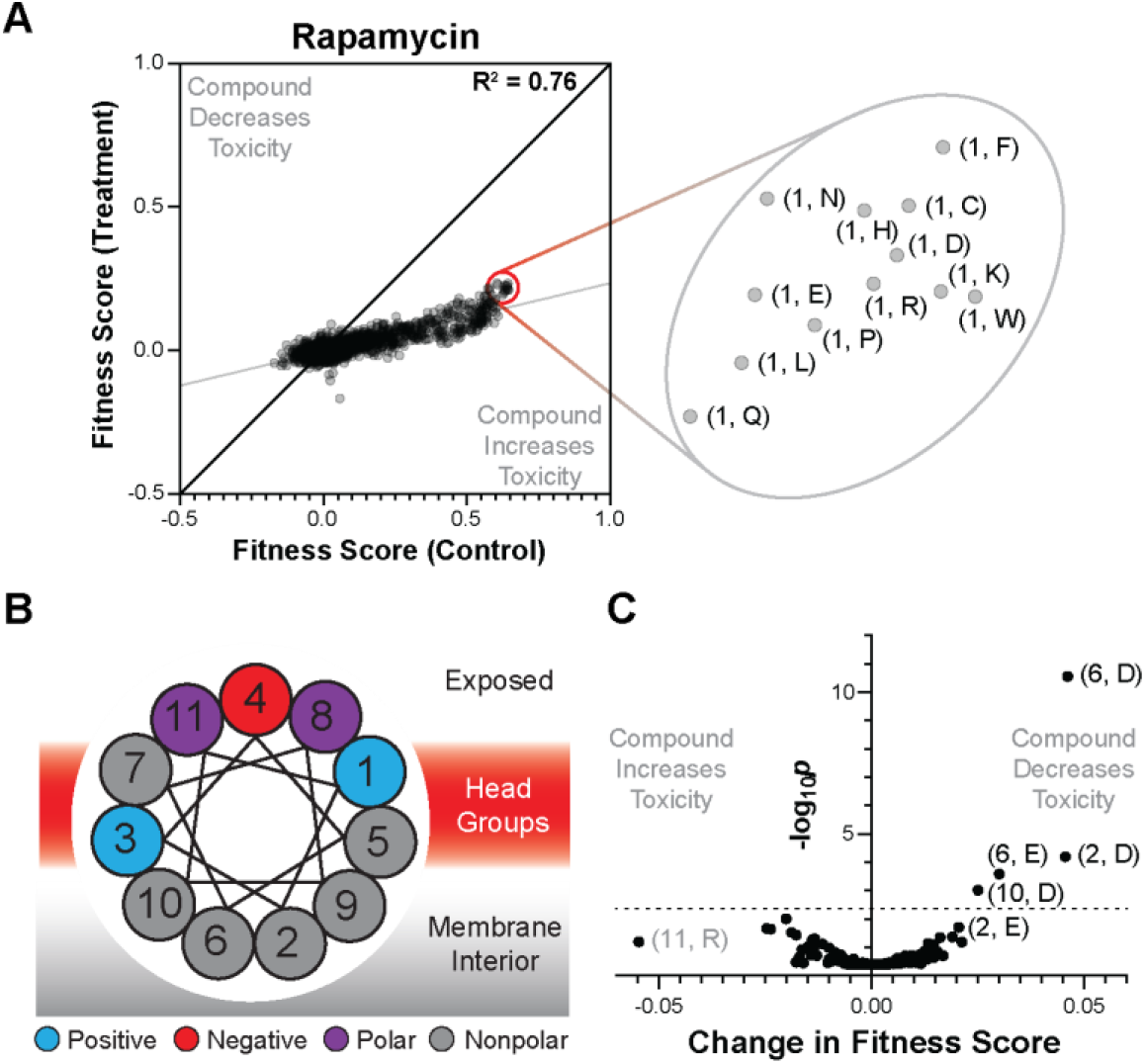
(A) Fitness scores of cells expressing α-synuclein mutants [labeled as (position, amino-acid substitution)] obtained in the presence of rapamycin relative to untreated controls. (B) Helical wheel of α-synuclein depicting the relative orientation of each residue in the 11-residue repeat. Schematic coloring reflects the physicochemical properties of the moiety/environment (blue: positively charged; red: negatively charged; purple: polar; grey: nonpolar). (C) Volcano plot describing changes in fitness score induced by rapamycin treatment, averaged over equivalent mutations in each of the seven repeating 11-residue segments, labeled as (position, amino-acid substitution). The significance of these changes was then determined from their z-scores, assuming a normal distribution, yielding the p-values shown. A 5% false discovery rate threshold is shown, which identifies the significance value above which 5% of the values would be falsely predicted as significantly different in the presence of compound; this threshold was determined by Benjamini-Hochberg method.

We next sought a model to explain the overall trend for variants with decreased affinity for membranes to have smaller effects in the presence of many of the compounds, and how this might be specifically modulated in the presence of rapamycin. We previously showed that α-synuclein mutants with reduced toxicity also had reduced affinity for binding to membranes.^6^ Moreover, the expression of α-synuclein in our system is tuned to maximize WT toxicity, so mutations that increase membrane affinity show little increase in toxicity. We hypothesize that a specific threshold of membrane-bound α-synuclein is necessary for toxicity, and that this threshold decreases under strong chemical stress.

Figure 6 relates the cellular protein concentration and dissociation constant to the fraction of binding; toxic binding regimes are indicated by a color gradient. This graph explains how fit mutants with reduced expression of α-synuclein can lead to decreased toxicity by lowering the overall surface coverage (Figure 6A). Similarly, mutations that decrease the affinity of α-synuclein for the membrane are expected to have the same effect (Figure 6A). Conversely, mutations predicted to increase membrane affinity, such as increases in the hydrophobicity of the membrane-binding face of the helix or the addition of positively charged residues in the interfacial region, have little effect on toxicity, likely because the toxic threshold for membrane binding by α-synuclein has already been exceeded (Figure 6A). If the fraction of binding required to achieve a toxic effect is decreased due to a pleiotropic chemical stress, then we see an expansion in the toxic region of the plot (Figure 6B), and a given suppressor will have a smaller effect on toxicity. Finally, treatment with rapamycin can be explained because it not only induces a pleiotropic effect on overall toxicity, but it also decreases the expression of the protein. Thus, the position of the WT shifts within the binding isotherm, resulting in a shift in the magnitude of the effect for the strongest suppressors (Figure 6C). Our data are therefore consistent with the hypothesis that α-synuclein exerts its toxicity in yeast through direct membrane interaction.

**Figure 6.**
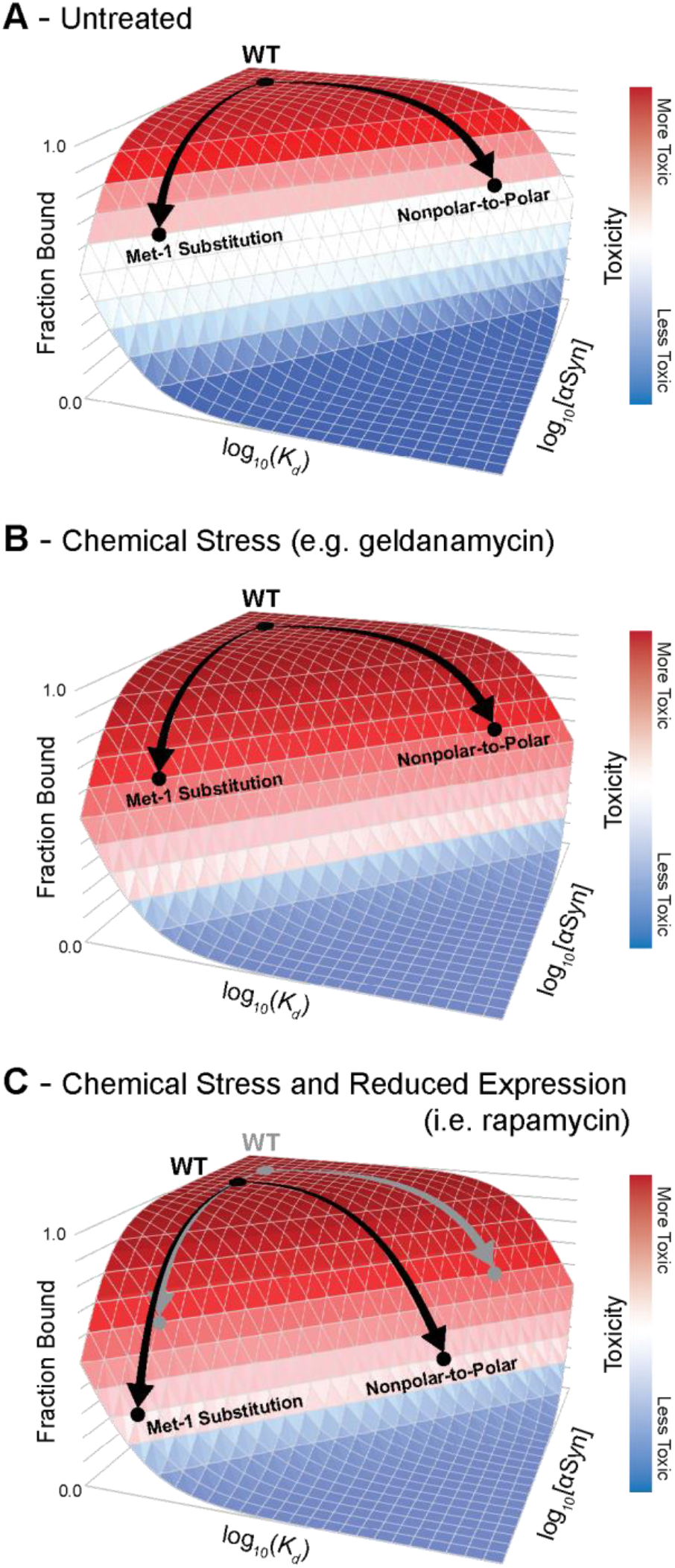
Cellular toxicity (represented by a red-white-blue color scale) depends on the membrane-bound population of α-synuclein. (A) In untreated cells, WT α-synuclein binds to cellular membranes above a critical threshold for toxicity. Mutations can reduce toxicity by decreasing the membrane-bound population of α-synuclein, either by decreasing membrane affinity (e.g., by introducing polar amino acids on the membrane-binding face) or reducing α-synuclein expression (e.g., by substituting Met-1). (B) Chemical stresses increase sensitivity to α-synuclein toxicity, so mutations that reduce toxicity in untreated cells are less effective at reducing toxicity under general chemical stress. (C) Because rapamycin reduces protein expression (black arrows relative to gray arrows), the combination of rapamycin and specific α-synuclein mutations reduces membrane binding sufficiently to overcome the increased sensitivity caused by compound toxicity.

This model is also supported by previous observations regarding the effect of genetic suppressors of α-synuclein toxicity in yeast. For example, overexpression of Rab proteins, which mediate the vesicle trafficking processes disrupted by α-synuclein, rescue α-synuclein toxicity only at moderate α-synuclein expression levels; no effect is reported at high α-synuclein concentrations.^57^ This is analogous to missense mutations, such as those in Figure 5C, that suppress α-synuclein toxicity under reduced expression conditions. In another example, the yeast gene YPP1 suppresses the toxicity of α-synuclein variants A30P, but not that of WT or A53T α-synuclein.^58^ The A30P mutant reduces membrane binding by disrupting formation of the amphiphilic helix, thereby reducing the population of the toxic conformation in yeast; only under this reduced α-synuclein burden is YPP1 able to suppress toxicity. Both these results and our own data suggest that α-synuclein can accumulate on cellular membranes at levels above that necessary to achieve toxicity, and therefore, many suppressors require a concomitant decrease in the population of the toxic species to demonstrate efficacy.

Comparing our results to other reports in the literature, there is also some evidence to suggest that a similar activity operates in neurons.^59^ For example, Dettmer, et al. report that mutations to the conserved repeat region, which modulate membrane binding, affect toxicity and inclusion formation in neuronal models.^60, 61^ Specifically, whereas WT α-synuclein is nontoxic under their expression conditions, substitutions predicted to increase membrane affinity drive inclusion formation and toxicity. Similar mutants were also found to have potent toxicity in animals.^62^ In neurons, it appears that mutations that enhance membrane affinity can increase the membrane-bound population of α-synuclein above a toxic threshold; in contrast, expression of α-synuclein in yeast under control of the GAL promoter likely causes the membrane-bound population to exceed a toxic threshold, which can be abrogated by mutations that disrupt membrane affinity. We speculate that dysregulation of membrane binding by α-synuclein could impart toxicity and reduce neuronal function by interfering with normal vesicular trafficking processes, including synaptic vesicle trafficking; indeed, yeast genes that rescue α-synuclein toxicity are enriched in factors that potentiate various trafficking processes.^20, 21^ Conversely, data from neither yeast nor neurons readily explains the toxicity of the PD-associated A30P mutant, which reduces membrane affinity, suggesting that additional pathological processes operate in humans.

### Membrane Binding Predicts Yeast Toxicity

In summary, chemical perturbation can affect the efficiency of disrupting mutations by changing the sensitivity of the cells to pleiotropic stress, while the pattern of mutations that alter toxicity remains unchanged. These findings provide strong support for the importance of a previously proposed structural model of α-synuclein, in which the N-terminal region serves as a membrane anchor, with increasing disorder and decreased membrane association along the chain. However, within this overall framework, there are two possible limiting possibilities that might account for the existing DMS and spectroscopic observations. In one model, the helix would become progressively less deeply submerged in the membrane and more dynamic toward the C terminus. A second limiting model would predict that any region of the chain could exist in an equilibrium between two distinct states; a *membrane-inserted helical state*, in which the helix is buried to a similar depth of the membrane at each position throughout the membrane-binding region, and a second, *non-inserted state* in which the chain is located outside the hydrophobic region of the membrane, adopting a largely random-coil conformation. In the two-state model, the decrease in the effect size of mutations along the chain would be associated with changes in the local preference for the membrane-inserted helical state (transitioning from strongly favoring the inserted state near the N terminus to favoring the non-inserted state near the C terminus).

We expected that these two models would have different sensitivities to mutations. Specifically, if the helix changes depth across its length, then different amino acids should be preferred at different positions. For example, Arg and Lys residues have a strong affinity for the headgroup region but are strongly disfavored within the interior of the bilayer; if the helix is well-buried at the N terminus and barely submerged at the C-terminus, the mutations to Arg or Lys on the membrane-bonding face should be unfavorable at the N terminus and favorable at the C terminus.

To address this possibility, we estimated the free energy required to transfer an amphiphilic helix with α-synuclein’s sequence to varying depths within the lipid bilayer. These depth-dependent energies can be estimated with significant precision based on the depth at which each amino acid is found in native membrane protein structures.^63^ If an amino acid requires greater energy to penetrate to a particular depth (e.g., positioning Asn at the center of the bilayer), then it will be rarely found at that depth in native membrane protein structures. Conversely, amino acids with favorable transfer energies (e.g., position Leu at the center of the bilayer) should be highly populated in native membrane protein structures. The observed depth-dependent propensities of each amino acid can be converted to pseudoenergies based on the Boltzmann relationship.

We used these pseudoenergies to estimate the binding energy of successive segments of α-synuclein to determine if decrease in mutational sensitivity results from a change in depth or a change in affinity. To do so, we constructed a structural model of an amphiphilic helix oriented parallel to the surface of an implicit lipid bilayer. The depth of each residue varied sinusoidally around an average depth of penetration; we set the periodicity of this helix to match that of the sequence–toxicity landscape (3.67 residues per turn)^6^ and set the phase of the helix to bury the hydrophobic residues to greatest depth. We calculated the transfer free energy associated with burying each residue to the expected depth based on depth-dependent pseudoenergies determined previously. We then repeated this calculation for a variety of depths of the helix axis (i.e., the average depth of penetration). To estimate the effect of changes in this transfer free energy caused by each missense mutation on the membrane-bound population of α-synuclein, we converted the estimated changes in transfer free energy to an expected change in bound fraction. This conversion requires an estimate of the transfer free energy of the WT protein, so we therefore repeated this computation for a wide variety of putative WT transfer free energies.

Our computational model therefore has two free parameters that allow us to discriminate between the two limiting conceptual models for the decrease in mutational sensitivity of α-synuclein toward the C terminus: average depth of penetration and transfer free energy of the WT protein. To determine the optimal values for these two parameters, we calculated the expected changes in the membrane-bound population for a range of values for each parameter and identified those that produced the highest correlation with the observed toxicity scores. To compare different regions of α-synuclein, we determined the optimal average depth of penetration and WT transfer free energy for successive segments, based on correlation with the experimental toxicity scores. α-Synuclein comprises seven repeated 11-residue segments that mediate membrane binding, so we determined the optimal parameters for each of these segments. Considering that over 200 data points are considered for each segment (20 mutations at each of 11 positions), the optimized parameters are well defined.

The model predicts that the average depth of insertion was nearly identical (13.9 ± 0.6 Å from the bilayer center) for each the 11-residue segment (Figure 7A), which is consistent with orthogonal measurements by EPR spectroscopy.^64^ To further rule out changes in depth as a driver of the decrease in mutational sensitivity toward the C terminus, we examined the correlation of our toxicity data with predicted changes in membrane-bound α-synuclein for a wide variety of depths. For mutations in each 11-residue membrane-binding segment, we predicted the change in membrane-bound α-synuclein that would result if that segment were to locally penetrate to depths between 10 and 20 Å from the bilayer center. We then calculated the Pearson correlation coefficient between those predicted changes in membrane-bound α-synuclein and the corresponding yeast toxicity data. For each segment, we found that changing the depth of penetration outside of a narrow, 1.5 Å window decreased the statistical correlation between the predictions of the model and the experimental data (Figure S11).

**Figure 7.**
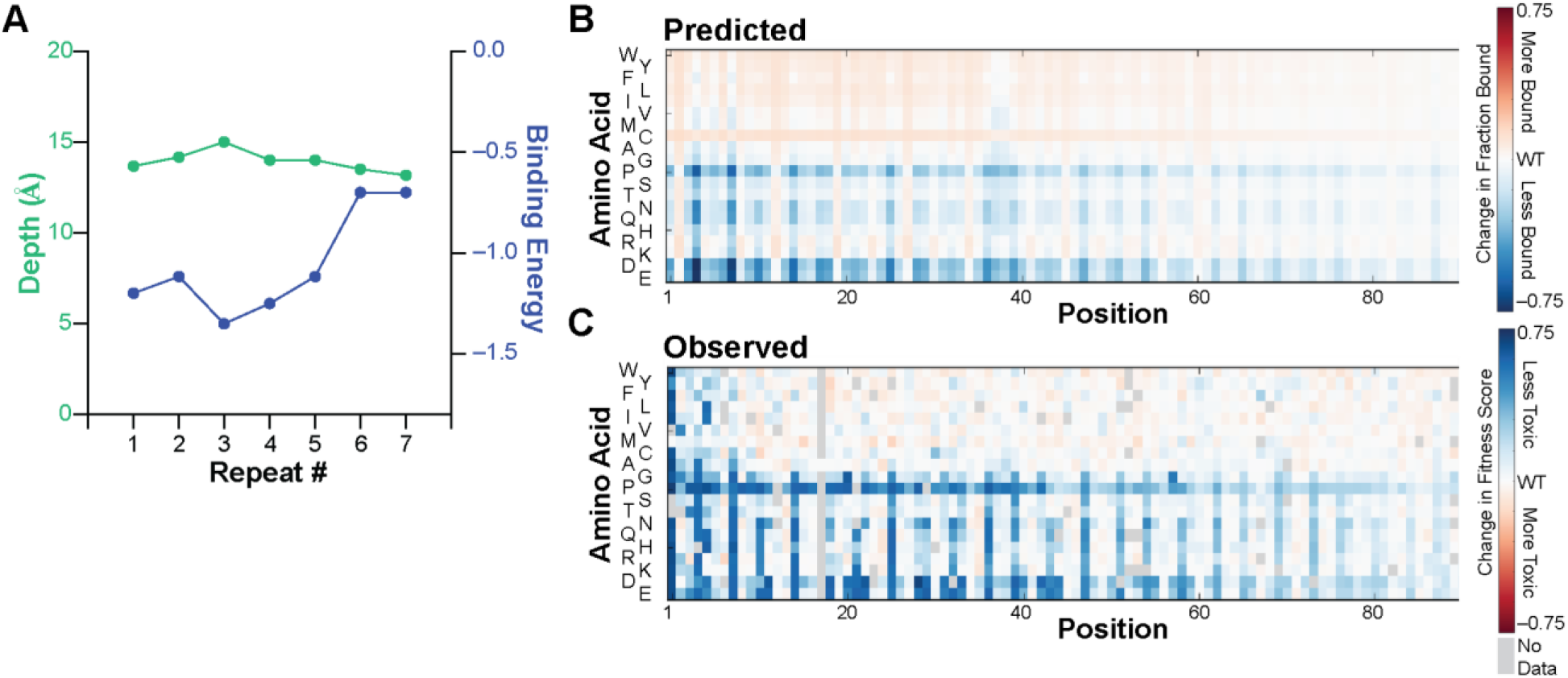
A thermodynamic model for α-synuclein–membrane interactions reveals determinants of decreasing mutational sensitivity. (A) The depth of insertion (green) and transfer free energy (blue) of each repeated 11-residue membrane-binding segment of α-synuclein were estimated by optimizing correlation between experimental toxicity scores and changes in the membrane-bound population of α-synuclein predicted based on the energy required to transfer an amphiphilic helix with the appropriate sequence to a particular depth within the lipid bilayer. See text for additional details. (B) Predicted changes in membrane-bound α-synuclein caused by missense mutations, based on the optimal penetration depth and transfer free energy of the WT protein determined in panel A and the depth-dependent transfer free energy of each amino acid. See text for additional details. Relative change in fraction bound is denoted as a red-white-blue color scale. (C) Observed changes in yeast toxicity (represented by a red-white-blue color scale) caused by α-synuclein missense mutations. The Pearson R^2^ correlation coefficient between all predicted and observed scores is 0.57.

Instead, we found that the predicted binding energy progressively decreased toward the C terminus (Figure 7A). Thus, the decrease in mutational sensitivity toward the C terminus likely results from reduced binding affinity rather than shallower insertion, resulting in partitioning between an ordered, membrane-bound helix and a disordered, dissociated coil, which is well-supported by spectroscopic studies.^23, 24^

Having established that decreases in mutational sensitivity correlate with computed changes in population of the bound helix and not from changes in depth, we sought to develop a model that would describe the observed sequence–toxicity landscape. We therefore predicted changes in the bound fraction of α-synuclein for the optimal depth and average WT transfer free energy determined previously (Figure 7A) and scaled the predicted changes in bound fraction by the observed decay in mutational sensitivity to account for decreases in the population of the membrane-bound helix toward the C terminus. The resulting predicted changes in the membrane-bound fraction agree remarkably with the observed fitness scores (Figure 7B/C), providing strong additional evidence for the membrane-bound helix model. In addition, the favorable binding energy predicted for the WT protein further support our conceptual model for the yeast toxicity of α-synuclein. Specifically, mutations that would be predicted to increase membrane binding (e.g., increases in the hydrophobicity of the membrane-binding face of the helix or the addition of positively charged residues in the interfacial region) have little effect on the membrane-bound population and therefore the cellular toxicity, whereas mutations that decrease membrane affinity (e.g., incorporating polar residues on the membrane-binding face of the helix) cause substantial changes in membrane binding and cellular toxicity.

This model is also consistent with the ability of membranes to induce α-synuclein amyloid formation.^65^ The disordered C terminus of the membrane-bound helix contains the aggregation-prone non-amyloid component (NAC) segment that forms the core of α-synuclein amyloids. Membrane binding increases local concentration of α-synuclein monomers, so if the aggregation-prone region releases from the membrane locally, it would be primed for aggregation.^66^

### Parallelization provides authentic research experience in the classroom

These studies were undertaken as an inquiry-based course for first-year biophysics and bioinformatics graduate students, which allowed the parallelized exploration of cellular mechanisms that mediate α-synuclein toxicity. In turn, these experiments provided training in experimental and computational skills, quantitative approaches to biological discovery and hypothesis testing, and key chemical biology concepts. Course-based research experiences have been demonstrated to promote the development of a wide variety of higher-order skills prioritized by leading scientific societies.^67^ To determine if our course was successful in achieving these goals, we performed a retrospective survey of course participants two to three years after course completion. Students were asked to rate their agreement with statements regarding the impacts of the course using a scale from 1 (strongly disagree) to 5 (strongly agree). With 89% of students responding, we found that students overwhelming agreed that the course improved their abilities across a number of areas (minimum average agreement of 4.0 on a 5-point scale; see below and Table S5 for complete results). The complete syllabus for the α-synuclein-focused course, which took place in two consecutive years, is available online: http://kampmannlab.ucsf.edu/pubs-2017 and http://kampmannlab.ucsf.edu/pubs-2018.

To adapt this study for the classroom, we divided each student cohort into teams of 3–4 with complementary academic backgrounds, ensuring computational and experimental expertise in each team, and assigned each group a teaching assistant who had completed the course previously. Approaching this project in teams proved one of the more valuable aspects of the course to students in retrospect; when surveyed two to three years after completing the course, students strongly agreed that the experience “improved [their] ability to work in interdisciplinary teams” (agreement 4.8 on a 5-point scale). This course met for three four-hour sessions per week for six weeks. Although the nature of yeast growth makes it advantageous to schedule several sessions in the same week, other biological systems (e.g., *E. coli*) would be amenable to other course schedules. In particular, quarter- or semester-long adaptations would provide opportunities to structure entire class periods around particular topics of focus (e.g., statistical approaches, instrumentation, etc.), which we instead covered organically as questions arose.

The first half of the course was devoted to data collection, with the remainder devoted to data analysis. In the experimental portion, each team selected a chemical perturbation from a curated list (Table 1). Teams then optimized treatment conditions, performed the library selection by outgrowth, and prepared DNA libraries for next-generation sequencing; in parallel, one of the instructors (R.W.N.) collected data in the absence of chemical treatment (Figure 1). Gratifyingly, these experimental exercises “improved [students’ self-reported] abilities in experimental laboratory work” (agreement 4.3 on a 5-point scale). During incubation periods, students formulated hypotheses for potential interactions of α-synuclein with the cellular processes perturbed by their compound. Students then wrote custom software in Python to process sequencing reads, build models, and test their hypotheses. During this process, instructors provided guidance in appropriate statistical approaches, computational tools, and visualization strategies. As a result, students reported that the course “improved [their] abilities in computational analysis” (agreement 4.5 on a 5-point scale) and “improved [their] ability to apply statistics to research questions” (agreement 4.0 on a 5-point scale). Throughout the course, students participated in a journal club covering papers relevant to the biological and technical foci of the course, which were supplemented by topical lectures by guest faculty. Teams also provided oral updates to their classmates, and a final presentation was given in an open symposium.

The generation, analysis, and interpretation of large datasets not only illustrated the powerful intersection of experiment and computation but also provided training in quantitative methods and reasoning, which are becoming increasingly essential for modern biological research. Students reported that the course “improved [their] ability to work with large datasets” (agreement 4.3 on a 5-point scale), suggesting that course like ours are an effective introduction. The focus on large genetic libraries, which have become readily available, provided ample opportunities for students to propose their own biological hypotheses at the data analysis stage and formulate quantitative, testable predictions, given the information content in these datasets. This approach is particularly suited to advanced training (e.g., upper-division or graduate-level), as it provides a more complex and open-ended opportunity than those typical of lower-level course-based research experiences, where hypotheses and analytical approaches are often prescribed. Indeed, students self-reported both “improved ability to propose and test hypotheses” (agreement 4.6 on a 5-point scale) and “improved ability to devise [one’s own] experiments/analyses” (agreement 4.2 on a 5-point scale), which were critical goals of our course.

Our approach also provided practical benefits for open-ended inquiry in laboratory courses. First, conducting experiments in pools reduced the impact of variation between students or teams, since the fitness of individual variants are calculated relative to the entire population as a whole. Each group’s experiment has WT as the internal control; thus, each group’s results can be internally normalized to facilitate comparison between groups. In this way, experimental or sampling errors (e.g., pipetting) are minimized. Second, by combining chemical perturbations with a single genetic variant library, we were able to address a variety of questions without the need for multiple libraries or other specialized biological materials. This allowed to us to provide each student or group of students with ownership of a complex project that relates to the broader goals of the course.

These particular experiments also illustrate many core biological concepts. Identifying the physical properties and chemical features of α-synuclein that mediate its activity illustrates the core concept of structure–function relationships. The DMS approach also offers a concrete example of the evolution of populations under various environmental constraints and illustrates gene–environment interactions that shape both normal and disease biology. Finally, by employing pleiotropic perturbations, we illustrate the interconnectedness of cellular systems. Students found this exposure valuable, reporting that the course “improved understanding of important biological concepts” (agreement 4.6 on a 5-point scale). We have focused on proteostasis, but other biological topics are readily addressed.

This framework was based on a previous version of our course, in which we examined environmental constraints on the fitness landscape of ubiquitin.^34, 35^ After thorough examination of the fitness of ubiquitin variants under many environmental stresses, we sought new subject matter to reinvigorate the course. Authentic, open-ended inquiry is central to achieving our learning goals of building independent critical thinking and fostering scientific creativity, which therefore required us to address previously unanswered questions. Without having to change the infrastructure of our course, which already featured yeast genetics and next-generation sequencing, we were able to sustain the course for more student cohorts. We therefore believe that this robust and adaptable approach provides a model for other graduate training experiences in classroom sizes typical of undergraduate laboratories.

## METHODS

### Overview of Library Construction (performed previously)^6^

In order to examine the relative toxicity of α-synuclein variants, we designed a barcoded DNA library to encode each of the 2,660 possible missense mutations of the protein. Double-stranded protein-coding DNA was produced commercially by solid-phase oligonucleotide synthesis and microfluidic assembly. This DNA library was cloned into a yeast vector under control of an inducible promoter. We then appended random 26-nucleotide barcodes downstream of the stop codon. These barcodes serve two purposes: (1) they simplify quantification of relative fitness by decreasing the necessary read length from 420 bp to less than 50 bp, which is easily accessible by Illumina sequencing, and (2) they provide a source of internal redundancy, since the fitness of each variant can be estimated from the change in frequency of ~11 independent barcodes.

### Library Cloning (performed previously)^6^

A pooled double-stranded DNA library containing the sequences of 2,660 α-synuclein mutants was generously provided by Twist Bioscience and cloned into pYES2-αSyn-EGFP, which was generously provided by the laboratory of V. M.-Y. Lee. Briefly, the α-synuclein library was amplified using primers specific to overhangs outside of the α-synuclein sequence. The product was purified by gel extraction and used as a megaprimer for amplification of the pYES-αSyn-EGFP plasmid by the EZClone method (Agilent). Following DpnI digestion, products were column purified and transformed into TG1 cells (Invitrogen) by electroporation, which produced 2.5M transformants. The transformed culture was grown overnight, and plasmids were extracted by miniprep. A GFP sequence containing an N-terminal flexible linker was amplified with primers that append a random 26 nt barcode immediately after the stop codon. The product was purified by gel extraction and used as a megaprimer for amplification of the plasmid extracted from transformed TG1 cells using the EZClone method. Transformation into TG1 cells by electroporation produced 10M transformants. The transformed culture was grown overnight, and plasmids were extracted by miniprep. The resulting plasmid pool was then transformed into DH5α cells to produce 60,000 transformants. The transformed culture was grown overnight, and plasmids were extracted by miniprep. The resulting plasmid pool was subjected to deep sequencing (see below) and transformed into W303 by the LiAc/PEG/ssDNA method,^68^ producing 21M transformants. Transformed cultures were grown overnight, back-diluted, and returned to log-phase growth. From this culture, glycerol stocks were prepared in SCD-Ura with 25% glycerol, each containing ~500M cells. Plating of these cultures after one freeze-thaw cycle yielded approximately 100M cells, ensuring appropriate representation of the library.

### Barcode Association (performed previously)^6^

To link α-synuclein variants to specific barcode sequences, plasmids isolated from DH5α cells were amplified by PCR to isolate the α-synuclein–GFP–barcode DNA sequence using primers that append Illumina adapter sequences. This product was gel purified and sequenced on an Illumina MiSeq using custom Read 1, Index Read 1, and Read 2 primers with 305, 20, and 205 cycles, respectively. Read 1 and Read 2 sequence α-synuclein and the index read sequenced the barcode.

Clusters with a greater than 5% chance of at least 1 error in the barcode sequence, judged by quality scores, were rejected from further analysis. Full-length α-synuclein sequences were constructed by merging paired-end reads. Non-overlapping regions were taken as the base call from the appropriate read. Bases in overlapping regions were taken as the consensus. In the event that the two reads disagreed, but one read indicated the WT base, the WT base was called. In the event that the two reads disagreed, and neither read indicated WT, the base call with the higher quality score was taken. Resulting merged reads were aligned against the synthesized sequences using Bowtie2, giving a set of barcode–variant pairs, one for each read. The list of variants mapping to each barcode was compiled across the entire set of reads, and accurate barcode– variant mappings were taken as those for which the most common mapping was at least twice as common as the next most common mapping. This workflow yielded a dictionary mapping 31,484 barcodes to 2,600 α-synuclein variants.

### Optimization of Selection Experiments

To identify appropriate concentrations of chemical treatments to perturb yeast cell biology, we measured the growth rate of yeast cells expressing WT α-synuclein in the presence of each compound over several orders of magnitude of concentration (Figure S12). Yeast cells were diluted to OD ~0.2 in SCD-Ura and grown for 12 hours with shaking at 30 °C. The final OD was then recorded and used to calculate a doubling time. Compound concentrations were selected to measurably retard growth without inducing overt toxicity. Concentrations employed in the selection experiments typically yielded 2–5 generations in the 12 hours between each timepoint.

### Yeast Library Selection

Glycerol stocks of a yeast library expressing 2,600 barcoded missense variants of α-synuclein (prepared as described previously^6^) were thawed into SCD-Ura (synthetic complete dextrose media lacking uracil) and shaken overnight at 30°C. Cells were collected, washed, and resuspended in SCR-Ura (synthetic complete raffinose media lacking uracil) and maintained in log phase for 12 hours. Cells were collected for t = 0 h timepoints by centrifugation and stored at –80 °C. The remaining culture was diluted to OD 0.2 in SCR-Ura with 1% galactose to induce protein expression and the appropriate concentration of compound; 0.1% proline and 3 × 10^−3^ % SDS were added to improve cellular permeability of brefeldin A (Figure S1).^69^ After 12 hours (2–5 generations), cells were collected and the culture was diluted to OD 0.2 in SCR-Ura with 1% galactose maintain log-phase growth. Final aliquots were collected after 12 more hours of growth. Each selection was performed in duplicate (Figure S13), starting with a new glycerol stock.

### Deep Sequencing

Plasmids were isolated by miniprep^70^ and amplified by PCR to isolate barcode DNA. A first round of PCR amplified the barcode region of the plasmid using primers that append a unique index for each replicate and time point and the standard index sequencing primer binding site. Products were gel purified and then amplified in a second PCR reaction to append Illumina adapter sequences. Products were gel purified, pooled, and sequenced on the HiSeq 4000 in SE50 mode using a custom Read 1 primer to sequence the barcode.

### Determination of Fitness Scores

Barcode sequences with high-confidence mappings to protein variants (see *Barcode Association* above) were counted at each time point. A representative barcode mapping to WT was then selected for comparison and normalization. This barcode was identified by determining the median change in log-transformed frequency over time amongst all barcodes mapping to WT; the barcode associated with the median change in log-transformed frequency over time was selected as the median WT barcode for comparison and normalization. Counts for barcodes mapping to mutants were then divided the count of median WT barcode at each timepoint to normalize each mutant to WT. These ratios were then log transformed to reflect the exponential growth of the yeast cultures. The log-transformed ratios for each barcode were fit to a line over the three time points. Slopes for each barcode were averaged across replicates, and the average slopes for synonymous barcodes were averaged to determine a fitness score and associated error for each α-synuclein variant.

### Changes in Fitness Score Caused by Chemical Perturbation

To account for differences in the number of generations between each experiment, we first determined a linear correlation between the fitness scores of the treated and untreated cells (Figure 4) using orthogonal distance regression, which integrates the error associated with each fitness score. The slope and intercept of these regressions were used to calculate an “expected” fitness score in each condition (i.e., the fitness score that would be expected if the only effect of chemical perturbation is a change in growth rate of the yeast cell without any functional interaction with α-synuclein). Deviations from this linear behavior (i.e., residuals) were then calculated for each mutant in each condition and reported as the changes in the fitness score of each mutant caused by chemical perturbation (e.g., Figure 5C, S1-10).

The average change in fitness at a given position within the 11-residue repeat was determined by averaging equivalent mutations (e.g., incorporation of glutamine at the second residue of each repeat) at each position and the associated error was propagated from the errors of the individual fitness scores. The significance of these changes was then determined from their z-scores, assuming a normal distribution, yielding the p-values shown in Figure 5C. A 5% false discovery rate threshold was determined by identifying the significance value above which 5% of the values would be falsely predicted as significantly different in the presence of compound; this threshold was determined by Benjamini-Hochberg method.

### Prediction of Mutational Effects

We modeled α-synuclein as a membrane-bound helix whose axis is parallel to the surface of the lipid bilayer. The depth each amino acid penetrates into the membrane was modeled as varying sinusoidally around the depth of the helix axis, with period 3.67 residues (obtained by fitting the periodicity of the fitness scores as described previously) and amplitude 3.3 Å, the approximate radius of Cβ atoms from the central axis of an α-helix:

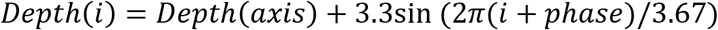

The phase of the helix was optimized empirically to position repeated hydrophobic residues at greatest depth. For varying depths of the helix axis, we calculated the energy associated with positioning each amino acid at the depth predicted from our structural model using energy potentials derived for each amino acid from their depth-dependent frequencies in high-resolution membrane protein structures.^63^ The energy difference between WT α-synuclein and each possible mutant (ΔΔG_mut_) was then used to predict a change in the membrane-bound fraction of α-synuclein resulting from substitution (FB_mut_).

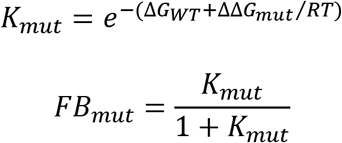

The average depth of penetration and transfer free energy of the WT protein were then optimized by grid search to provide the best linear correlation of predicted changes in bound fraction with observed changes in fitness score for each 11-residue membrane binding repeat (Figure 7A). The average optimal depth and WT transfer free energy across all 7 segments was then used to predict the change in bound population for all mutants (Figure 7B/C). To describe the observed attenuation in mutational sensitivity, we fit the fitness scores for residues, averaged at each position over a sliding 11-residue window, to a logistic equation, which captured the overall trend in the data. The product of the depth-dependent energies and this logistic scaling factor produce the predictions of effect size reported in Figure 7C.

### Retrospective Survey of Course Effectiveness

Students who completed the course were asked to complete an anonymous survey to gauge the success with which course goals were met; survey questions and results are listed in Table S5. Students were informed that the results might be published, but that respondent identities would remain anonymous. Students were further informed that neither completion of the survey nor any responses therein would impact their course grade, status in their program, or their status as a co-author of any relevant publications. Surveys were administered in July 2020, nearly two or three years after completion of the course by the Fall 2018 and Fall 2017 cohorts, respectively. Students were given four days to complete the survey, which resulted in an 89% response rate (32 students responding out of 36 total). Publication of student course evaluations was approved by the UCSF Institutional Review Board.

## Acknowledgments

Publication of student course evaluations was approved by the UCSF Institutional Review Board. We thank R.H. Edwards, S. N. Floor, J. E. Gestwicki, C. Joseph, T. Kortemme, D. D. Larsen, H. D. Madhani, and L. McConlogue for guest lectures in the course, J. S. Fraser for advice, the lab of H. El-Samad for yeast strains, the lab of V. M-Y. Lee for plasmids, and Twist Bioscience for providing the dsDNA variant library in support of our educational efforts. This work was supported by grants from the NIH to M.K. (DP2-GM119139) and to W.F.D. (R35-GM122603, P01-AG002132, R01-GM117593) and the UCSF Program in Breakthrough Biomedical Research, which is funded in part by the Sandler Foundation (Postdoctoral Fellowship to R.W.N., and award to M.K.). R.W.N. was supported by NIH grant T32-HL007731. T.A., G.C.H., A.M.M., T.S.W., T.B.C., Y.K.G., M.C.J., J.P., C.A.S., and S.A.W. were supported by NSF Graduate Research Fellowships. T.A., A.M.M., D.M.C.S., D.R.W., and T.S.W. were supported by NIH grant T32-GM064337. D.M.C.S. was supported by NIH grants F31-CA232325 and U54-CA209891. D.R.W. was supported by NIH grant F31-CA243439. T.B.C., M.G.J., and K.E.L. were supported by UCSF Discovery Fellowships. M.G.G. was supported by NIH grant F31-HG011007. Y.K.G. and M.C.J. were supported by NIH grant R25-GM056847.

## Author Contributions

R.W.N. and M.K. conceived the project and designed the course curriculum. R.W.N., E.D.C., and M.K. designed the library and sequencing strategy. R.W.N. constructed the variant library and yeast strains. R.W.N., T.A., J.C., G.C.H., A.M.M., Z.Z.C.N., M.P., N.R.R., D.M.C.S., D.R.W., T.S.W., D.B., C.C., A.C., T.B.C, L.S.E., B.F., E.A.F., M.A.G., Y.K.G., M.G.G., L.M.G., N.H., M.J.-G., M.C.J., M.G.J., A.F.K., K.E.L, J.L, C.M., E.E.M., L.E.M.-V., E.J.N., A.P., J.P., W.S., C.A.S., J.S., H.T., S.A.W., D.R.W., G.W., S.R., and E.D.C. contributed to data collection. T.A., J.C., G.C.H., A.M.M., Z.Z.C.N., M.P., N.R.R., D.M.C.S., D.R.W., and T.S.W. served as teaching assistants and provided guidance to student teams. R.W.N. and W.F.D. developed the structural model. All authors contributed to data analysis and writing of the manuscript.

## Supporting Information

Expanded methods; complete sequence–toxicity landscapes (Figures S1–S10); correlation of predicted and observed mutational effects for varying depths of membrane permeation (Figure S11); effects of compound treatments on yeast growth rate (Figure S12); inter-replicate correlations (Figure S13); table of complete fitness scores (Table S1); list of fitness scores and corresponding errors (Table S2); counts of each variant at each time point (Table S3); changes in fitness scores for variants more toxic than WT (Table S4); DNA sequences (Table S5)

## Data Availability

Barcode counts in each condition at each time point for each replicate are available as supporting tables, as are the resulting fitness scores and associated errors. Fitness scores are also available at MaveDB (mavedb.org; accession numbers mavedb:00000045-a-1 through mavedb:00000045-l-1). Raw sequencing data are available at the NCBI Sequence Read Archive (PRJNA564806). Custom software developed for data processing is available at github.com/rnewberry17.

## TOC GRAPHIC

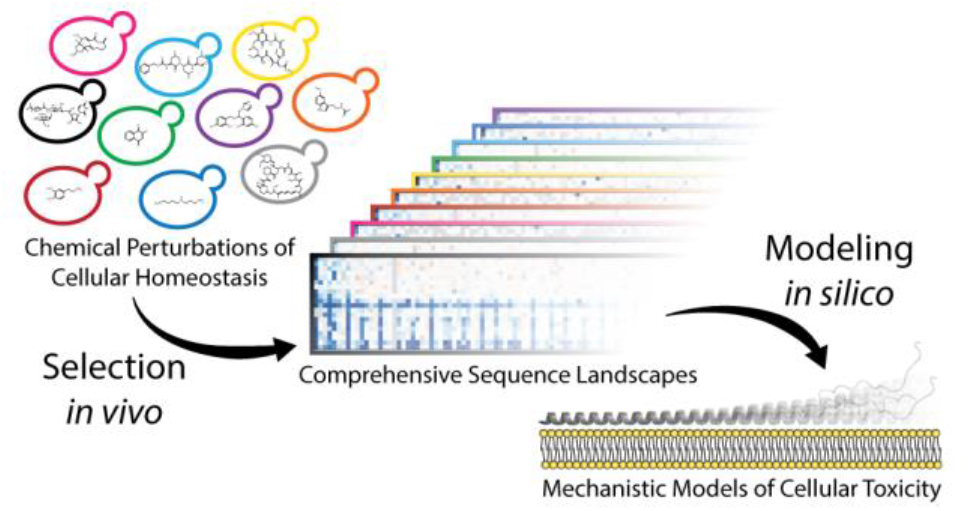

