## Supporting Information for "Robust Sequence Determinants of α-Synuclein Toxicity in Yeast Implicate Membrane Binding"

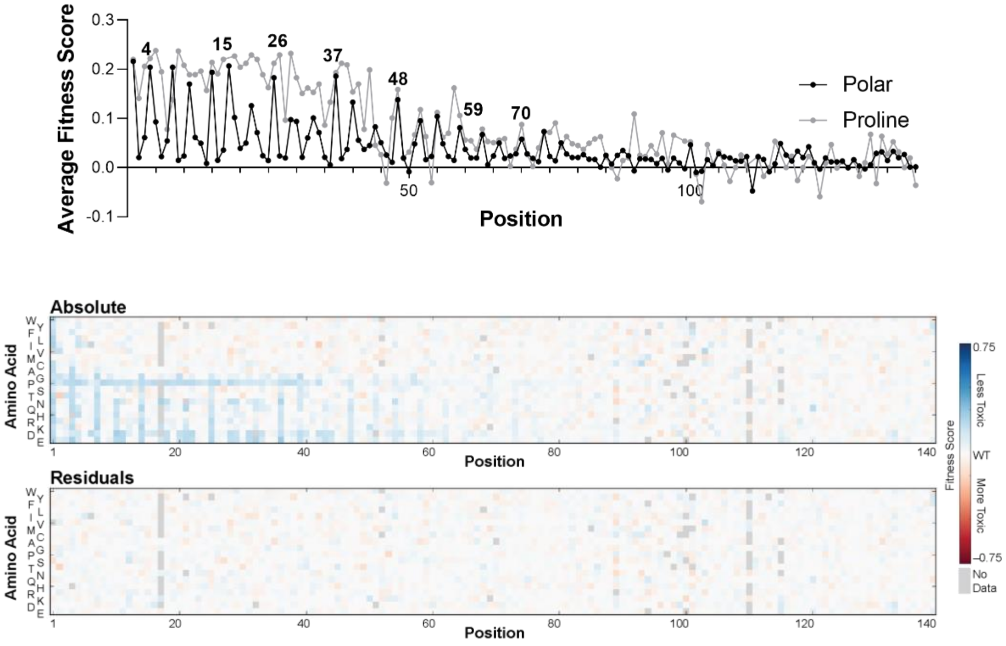


**Figure S1.** Changes in fitness scores of α-synuclein variants due to treatment of yeast with 40 µg/mL brefeldin A, 0.1% proline, 3 × $10^{-3}$ % SDS, relative to yeast treated with 0.1% proline, 3 × $10^{-3}$ % SDS.

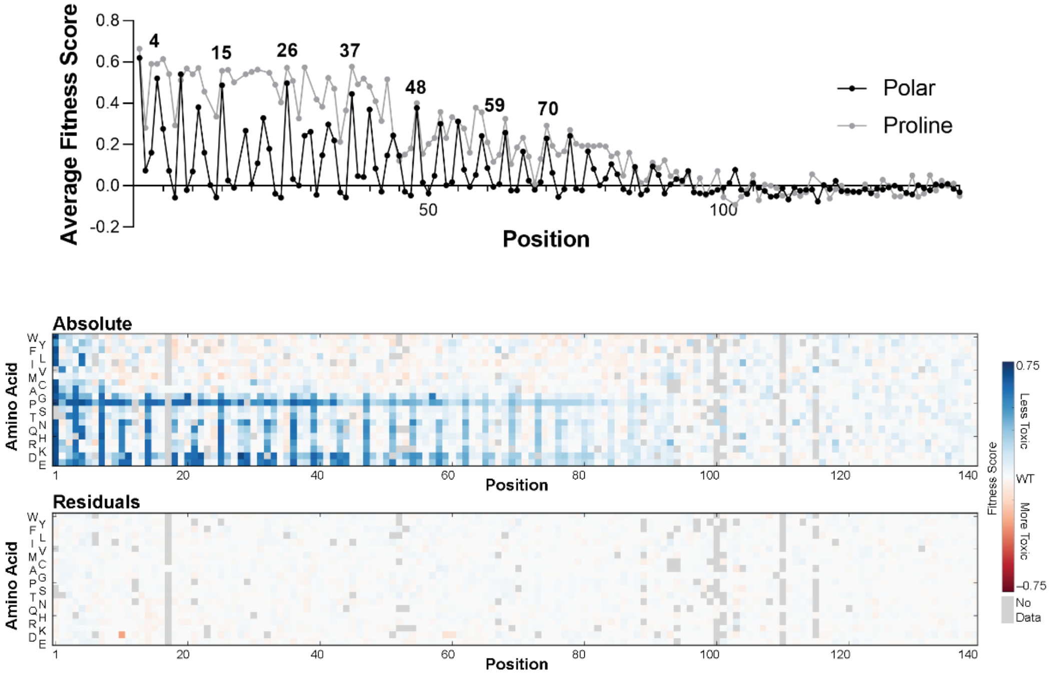


**Figure S2.** Changes in fitness scores of α-synuclein variants due to treatment of yeast with 20 mM dopamine.

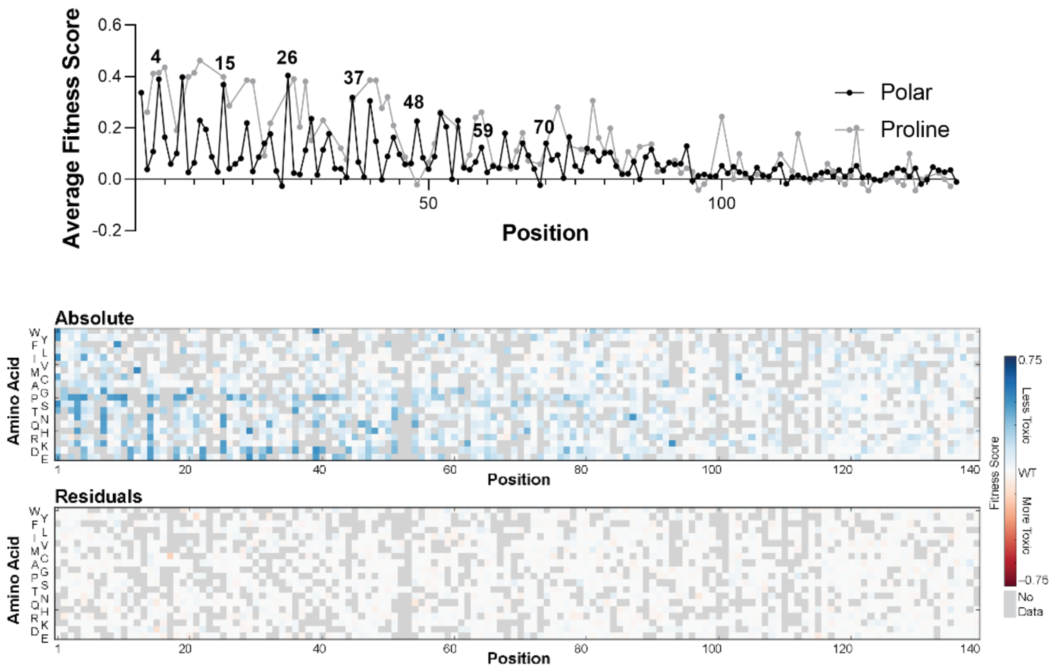


**Figure S3.** Changes in fitness scores of α-synuclein variants due to treatment of yeast with 20 μM geldanamycin.

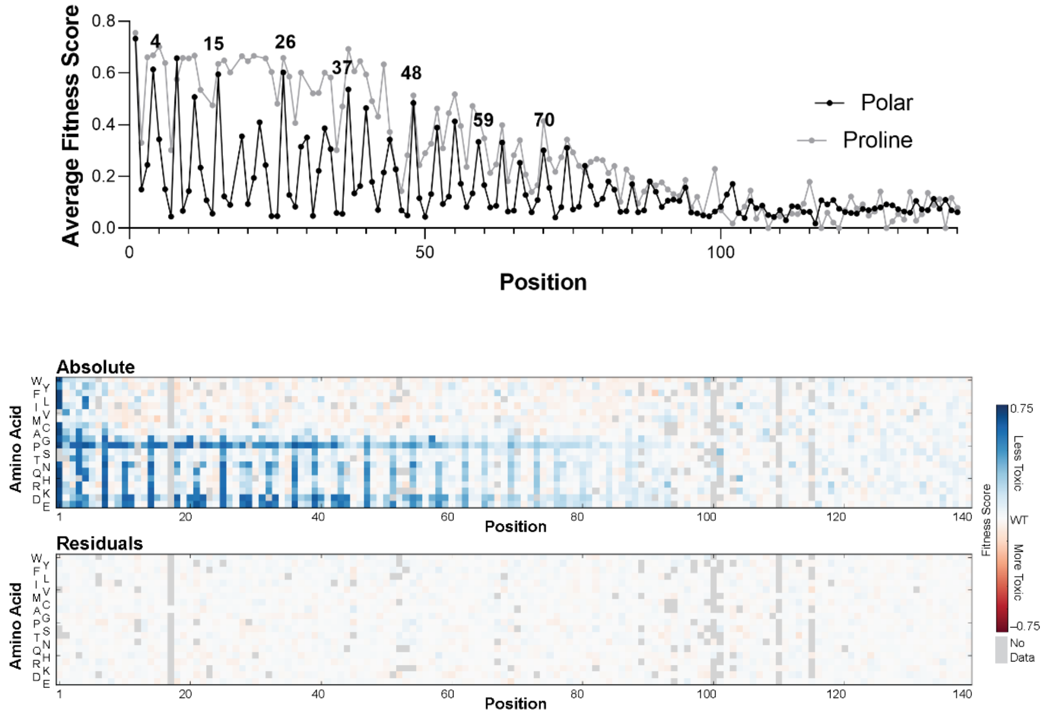


**Figure S4.** Changes in fitness scores of α-synuclein variants due to treatment of yeast with 250 μM melatonin.

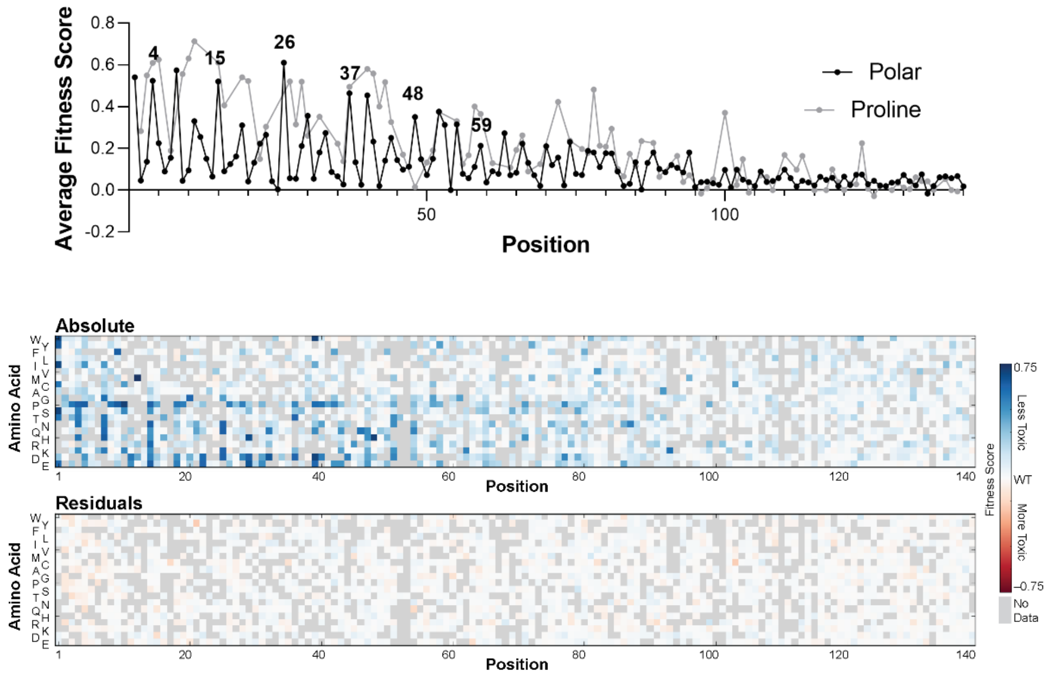


**Figure S5.** Changes in fitness scores of α-synuclein variants due to treatment of yeast with 40 µM menadione.

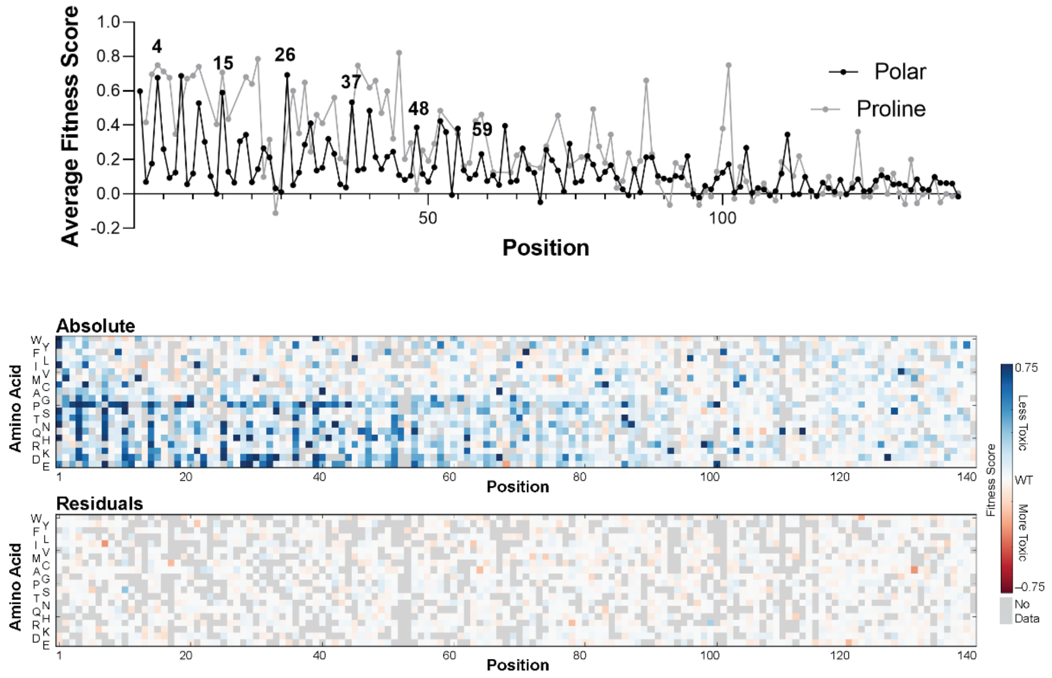


**Figure S6.** Changes in fitness scores of α-synuclein variants due to treatment of yeast with 130 µM MG-132.

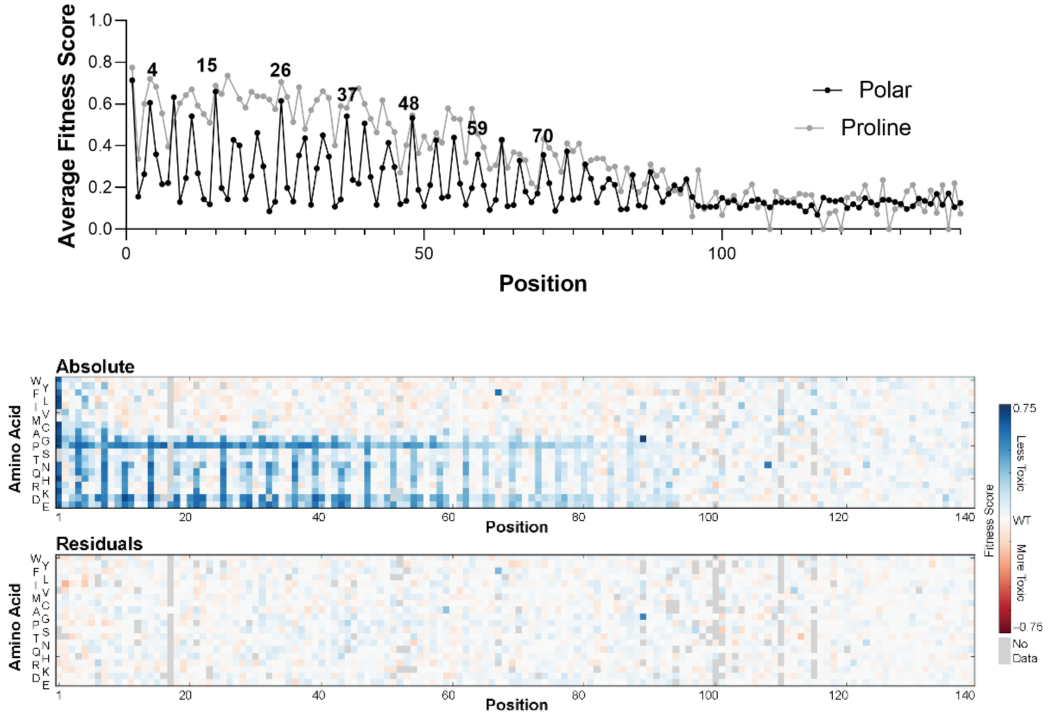


**Figure S7.** Changes in fitness scores of α-synuclein variants due to treatment of yeast with 1 µM miconazole.

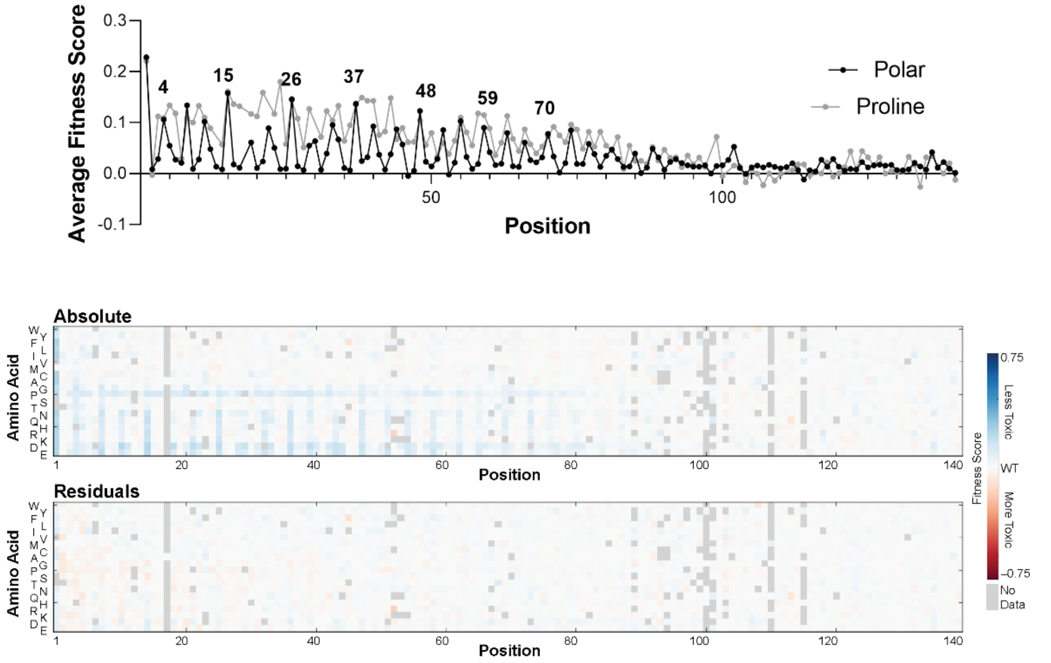


**Figure S8.** Changes in fitness scores of α-synuclein variants due to treatment of yeast with 10 ng/mL rapamycin.

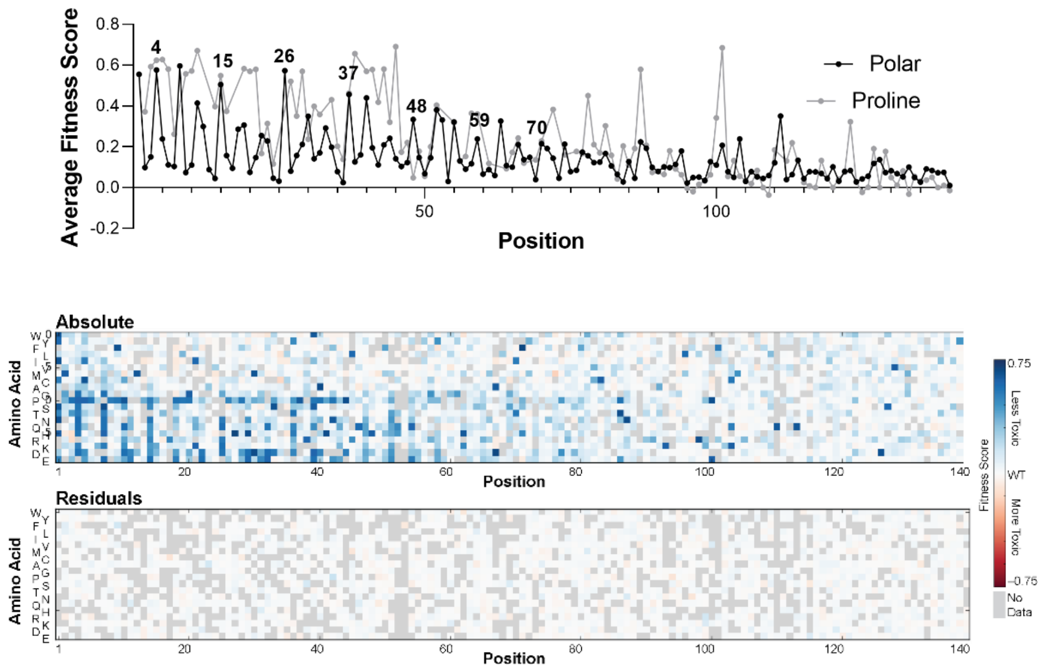


**Figure S9.** Changes in fitness scores of α-synuclein variants due to treatment of yeast with 1 mM spermidine.

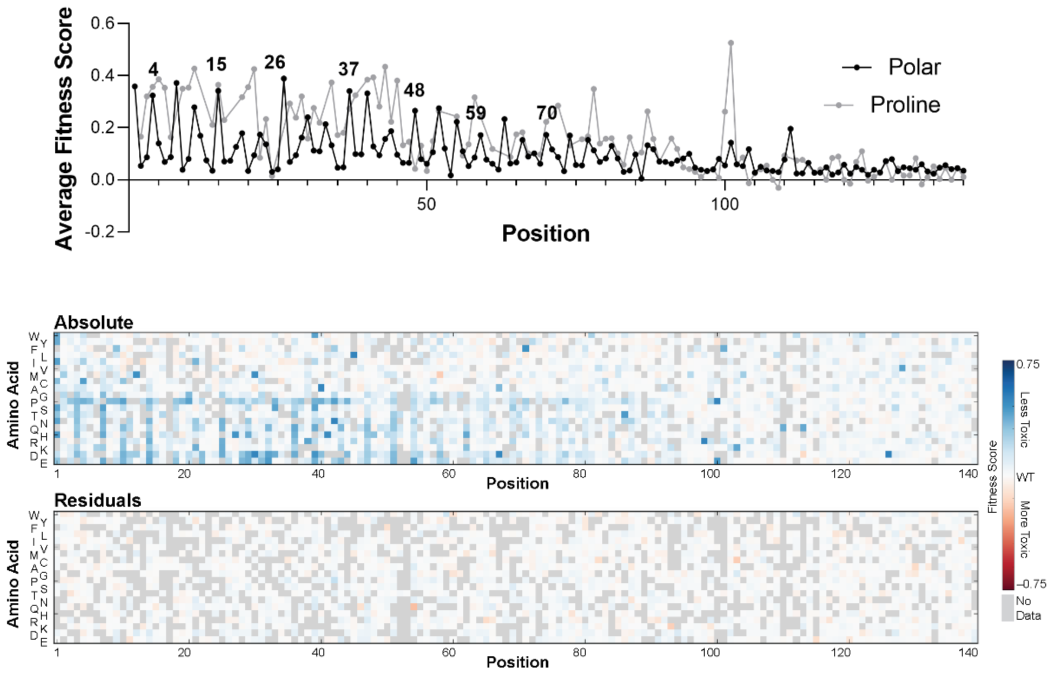


**Figure S10.** Changes in fitness scores of α-synuclein variants due to treatment of yeast with 100 ng/mL tunicamycin.

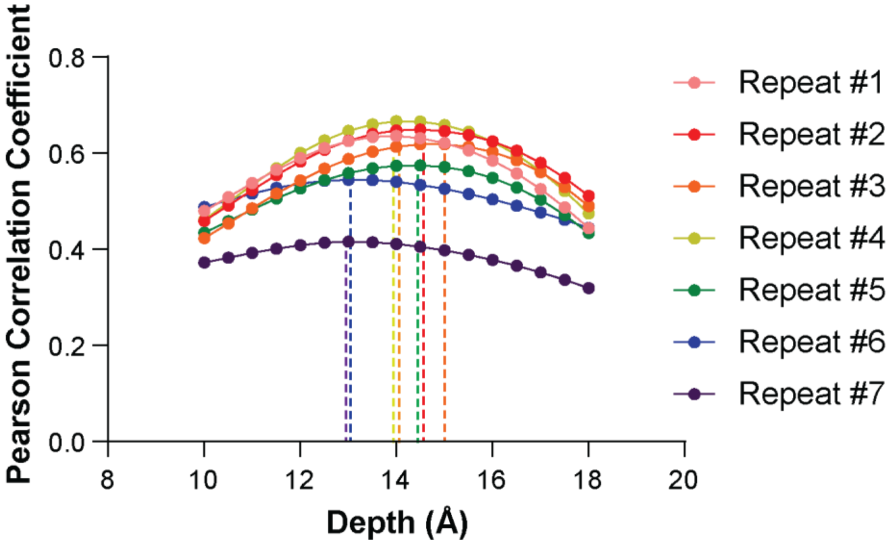


**Figure S11.** Correlation of experimental changes in α-synuclein toxicity with expected changes in membrane-bound α-synuclein predicted by structural modeling. For mutations within each 11-residue membrane-binding repeat segment, we predicted the change in membrane-bound α-synuclein, based on a structural model that embeds an amphiphilic helix to varying depths within a lipid bilayer (see "Prediction of Mutation Effects" above). We predicted the effect of each mutation for a variety of depths of penetration and then calculated the correlation of those predictions with the measured toxicity scores. For each 11-residue segment, we find that the Pearson correlation coefficient is maximized within a narrow, 1.5 Å window. Furthermore, changes in depth outside that window compromise the quality of the correlation. Therefore, we find less statistical support for varying depths of penetration than we do for a consistent depth.

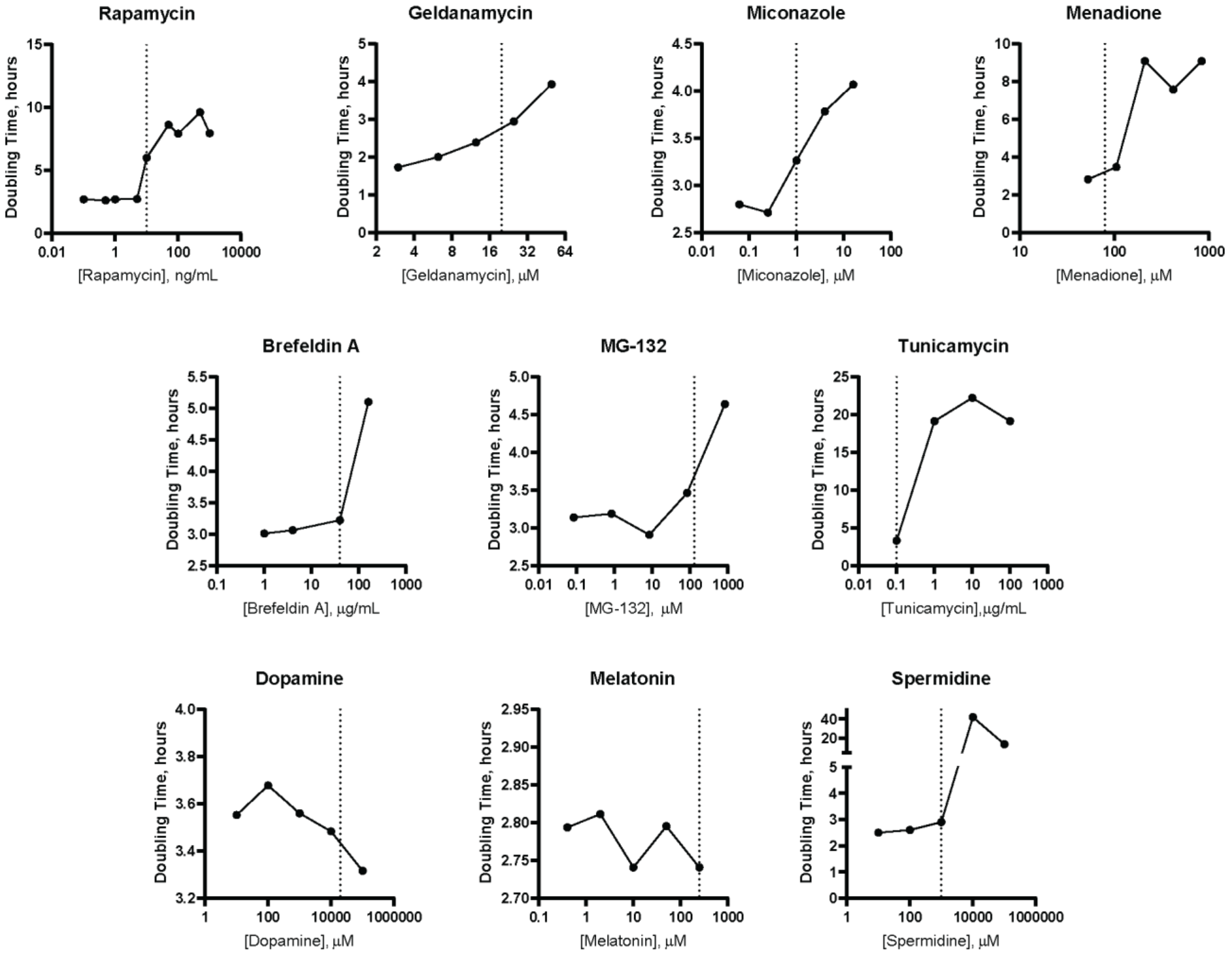


**Figure S12.** Changes in growth rate of yeast expressing WT α-synuclein when treated with varying concentrations of chemical perturbants. The concentrations selected for screening the deep mutational scanning library are shown as dotted vertical lines.

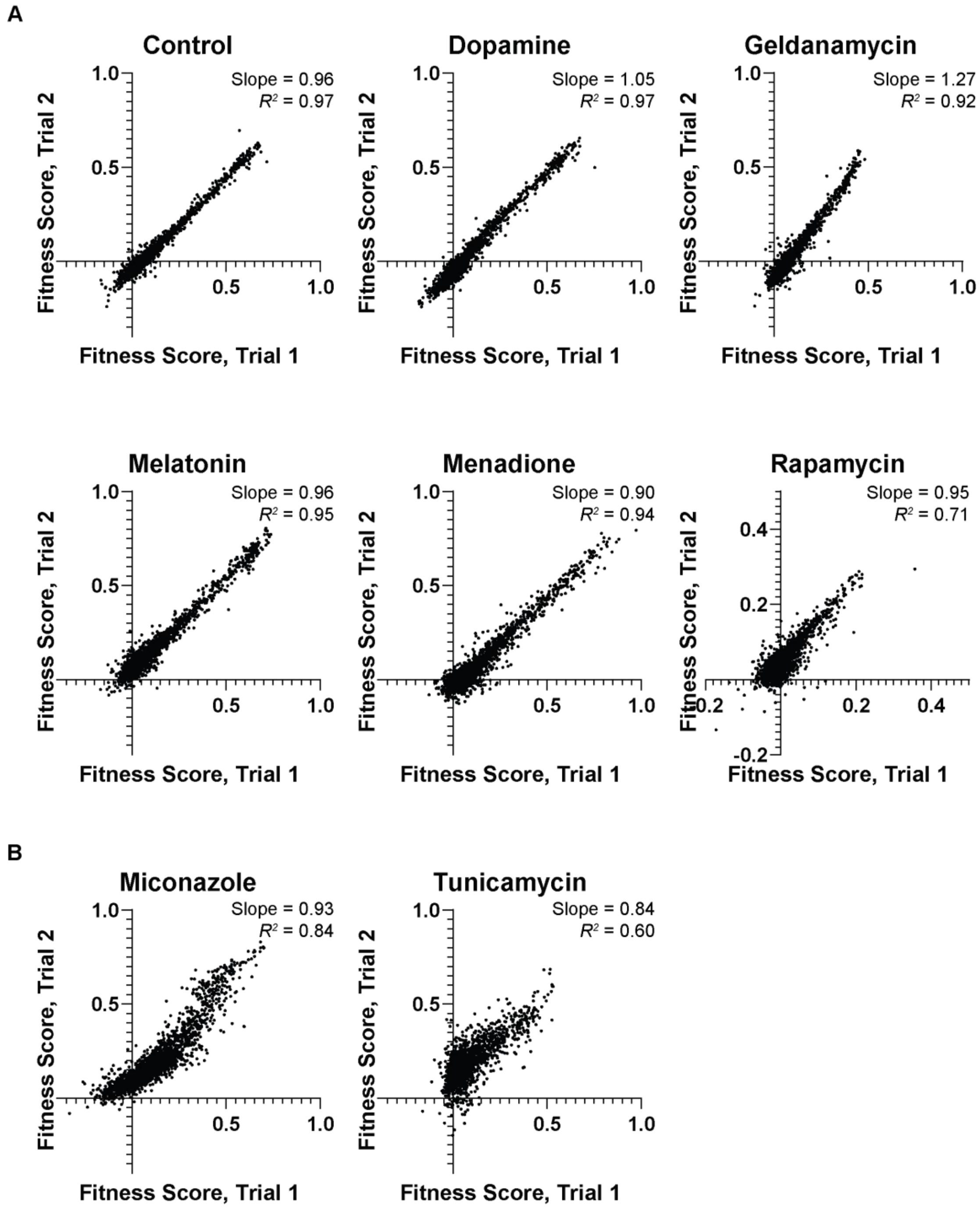


**Figure S13.** Correlation in fitness scores between replicates of selections performed in the presence of various small molecules. (A) Inter-replicate correlation for conditions with two 3-timepoint replicates. (B) For miconazole and tunicamyin, one replicate consisted of only two timepoints. We show the correlation between fitness scores obtained for both replicates to emphasize the reproducibility of the experiment; however, to allow comparisons with other treatments, we subject only the 3-timepoint replicate to further analyis. A single replicate was collected in the presence of brefeldin A, spermidine, and MG-132. All correlation coefficients are Pearson correlation coefficients.

**Table S5.** Results of retrospective survey on outcomes of UCSF course Biophysics 205A – Physical Underpinnings of Biological Systems (PUBS).

| | Number of Responses | | | | | |
|---|---|---|---|---|---|---|
| Statement | (1) Strongly Disagree | (2) Somewhat Disagree | (3) Neutral | (4) Somewhat Agree | (5) Strongly Agree | Average |
| PUBS improved my understanding of important biological concepts. | 0 | 0 | 1 | 12 | 19 | 4.56 |
| PUBS improved my ability to propose and test hypotheses. | 0 | 0 | 0 | 12 | 20 | 4.63 |
| PUBS improved my ability to devise my own experiments/analyses. | 0 | 1 | 6 | 10 | 15 | 4.22 |
| PUBS improved my abilities in experimental laboratory work. | 0 | 0 | 7 | 10 | 15 | 4.25 |
| PUBS improved my abilities in computational analysis. | 0 | 0 | 5 | 7 | 20 | 4.47 |
| PUBS improved my ability to apply statistics to research questions. | 0 | 2 | 6 | 13 | 11 | 4.03 |
| PUBS improved my ability to work with large datasets. | 0 | 0 | 6 | 10 | 16 | 4.31 |
| PUBS improved my ability to work in interdisciplinary teams. | 0 | 0 | 0 | 5 | 27 | 4.84 |

**Table S6.** DNA Sequences

| | |
|---|---|
| PCR 1 Forward Primer - Index 1 | GAAGAGCACACGTCTGAACTCCAGTCACCGTGATCGATCACATGGTCCTGCTGGAG |
| PCR 1 Forward Primer - Index 2 | GAAGAGCACACGTCTGAACTCCAGTCACACATCGCGATCACATGGTCCTGCTGGAG |
| PCR 1 Forward Primer - Index 3 | GAAGAGCACACGTCTGAACTCCAGTCACGCCTAACGATCACATGGTCCTGCTGGAG |
| PCR 1 Forward Primer - Index 4 | GAAGAGCACACGTCTGAACTCCAGTCACTGGTCACGATCACATGGTCCTGCTGGAG |
| PCR 1 Forward Primer - Index 5 | GAAGAGCACACGTCTGAACTCCAGTCACCACTGTCGATCACATGGTCCTGCTGGAG |
| PCR 1 Forward Primer - Index 6 | GAAGAGCACACGTCTGAACTCCAGTCACATTGGCCGATCACATGGTCCTGCTGGAG |
| PCR 1 Forward Primer - Index 7 | GAAGAGCACACGTCTGAACTCCAGTCACGATCTGCGATCACATGGTCCTGCTGGAG |
| PCR 1 Forward Primer - Index 8 | GAAGAGCACACGTCTGAACTCCAGTCACTCAAGTCGATCACATGGTCCTGCTGGAG |
| PCR 1 Forward Primer - Index 9 | GAAGAGCACACGTCTGAACTCCAGTCACCTGATCCGATCACATGGTCCTGCTGGAG |
| PCR 1 Forward Primer - Index 10 | GAAGAGCACACGTCTGAACTCCAGTCACAAGCTACGATCACATGGTCCTGCTGGAG |
| PCR 1 Forward Primer - Index 11 | GAAGAGCACACGTCTGAACTCCAGTCACGTAGCCCGATCACATGGTCCTGCTGGAG |
| PCR 1 Forward Primer - Index 12 | GAAGAGCACACGTCTGAACTCCAGTCACTACAAGCGATCACATGGTCCTGCTGGAG |
| PCR 1 Forward Primer - Index 13 | GAAGAGCACACGTCTGAACTCCAGTCACTTGACTCGATCACATGGTCCTGCTGGAG |
| PCR 1 Forward Primer - Index 14 | GAAGAGCACACGTCTGAACTCCAGTCACGGAACTCGATCACATGGTCCTGCTGGAG |
| PCR 1 Forward Primer - Index 15 | GAAGAGCACACGTCTGAACTCCAGTCACTGACATCGATCACATGGTCCTGCTGGAG |
| PCR 1 Forward Primer - Index 16 | GAAGAGCACACGTCTGAACTCCAGTCACGGACGGCGATCACATGGTCCTGCTGGAG |
| PCR 1 Forward Primer - Index 17 | GAAGAGCACACGTCTGAACTCCAGTCACCTCTACCGATCACATGGTCCTGCTGGAG |
| PCR 1 Forward Primer - Index 18 | GAAGAGCACACGTCTGAACTCCAGTCACGCGGACCGATCACATGGTCCTGCTGGAG |
| PCR 1 Forward Primer - Index 19 | GAAGAGCACACGTCTGAACTCCAGTCACTTTCACCGATCACATGGTCCTGCTGGAG |
| PCR 1 Forward Primer - Index 20 | GAAGAGCACACGTCTGAACTCCAGTCACGGCCACCGATCACATGGTCCTGCTGGAG |
| PCR 1 Forward Primer - Index 21 | GAAGAGCACACGTCTGAACTCCAGTCACCGAAACCGATCACATGGTCCTGCTGGAG |
| PCR 1 Forward Primer - Index 22 | GAAGAGCACACGTCTGAACTCCAGTCACCGTACGCGATCACATGGTCCTGCTGGAG |
| PCR 1 Forward Primer - Index 23 | GAAGAGCACACGTCTGAACTCCAGTCACCCACTCCGATCACATGGTCCTGCTGGAG |
| PCR 1 Forward Primer - Index 24 | GAAGAGCACACGTCTGAACTCCAGTCACGCTACCCGATCACATGGTCCTGCTGGAG |
| PCR 1 Forward Primer - Index 25 | GAAGAGCACACGTCTGAACTCCAGTCACATCAGTCGATCACATGGTCCTGCTGGAG |
| PCR 1 Forward Primer - Index 26 | GAAGAGCACACGTCTGAACTCCAGTCACGCTCATCGATCACATGGTCCTGCTGGAG |
| PCR 1 Forward Primer - Index 27 | GAAGAGCACACGTCTGAACTCCAGTCACAGGAATCGATCACATGGTCCTGCTGGAG |
| PCR 1 Forward Primer - Index 28 | GAAGAGCACACGTCTGAACTCCAGTCACCTTTTGCGATCACATGGTCCTGCTGGAG |
| PCR 1 Forward Primer - Index 29 | GAAGAGCACACGTCTGAACTCCAGTCACTAGTTGCGATCACATGGTCCTGCTGGAG |
| PCR 1 Forward Primer - Index 30 | GAAGAGCACACGTCTGAACTCCAGTCACCCGGTGCGATCACATGGTCCTGCTGGAG |
| PCR 1 Forward Primer - Index 31 | GAAGAGCACACGTCTGAACTCCAGTCACATCGTGCGATCACATGGTCCTGCTGGAG |
| PCR 1 Forward Primer - Index 32 | GAAGAGCACACGTCTGAACTCCAGTCACTGAGTGCGATCACATGGTCCTGCTGGAG |
| PCR 1 Forward Primer - Index 33 | GAAGAGCACACGTCTGAACTCCAGTCACCGCCTGCGATCACATGGTCCTGCTGGAG |
| PCR 1 Forward Primer - Index 34 | GAAGAGCACACGTCTGAACTCCAGTCACGCCATGCGATCACATGGTCCTGCTGGAG |
| PCR 1 Forward Primer - Index 35 | GAAGAGCACACGTCTGAACTCCAGTCACAAAATGCGATCACATGGTCCTGCTGGAG |
| PCR 1 Forward Primer - Index 36 | GAAGAGCACACGTCTGAACTCCAGTCACTGTTGGCGATCACATGGTCCTGCTGGAG |
| PCR 1 Reverse Primer | GGTTAGAGCGGATGTGGGGG |
| PCR 2 Forward Primer | AATGATACGGCGACCACCGAGATCTACACGATCGGAAGAGCACACGTCTGAACTCCAGTC |
| PCR 2 Reverse Primer | CAAGCAGAAGACGGCATACGAGATGGTTAGAGCGGATGTGGGGG |
| Custom Illumina Read 1 Primer | GGGATTACACATGGCATGGACGAGCTGTACAAGTAA |
| Plasmid Sequence | CTGCATTAATGAATCGGCCAACGCGCGGGGAGAGGCGGTTTGCGTATTGGGCGCTCTTCCGCTTCCTCGCTCACTGACTC<br>GCTGCGCTCGGTCGTTCGGCTGCGGCGAGCGGTATCAGCTCACTCAAAGGCGGTAATACGGTTATCCACAGAATCAGGGG<br>ATAACGCAGGAAAGAACATGTGAGCAAAAGGCCAGCAAAAGCCCAGGAACCGTAAAAAGGCCGCGTTGCTGGCGTTTTTC<br>CATAGGCTCCGCCCCCCTGACGAGCATCACAAAAATCGACGCTCAAGTCAGAGGTGGCGAAACCCGACAGGACTATAAAG<br>ATACCAGGCGTTTCCCCCTGGAAGCTCCCTCGTGCGCTCTCCTGTTCCGACCCTGCCGCTTACCGGATACCTGTCCGCCT<br>TTCTCCCTTCGGGAAGCGTGGCGCTTTCTCATAGCTCACGCTGTAGGTATCTCAGTTCGGTGTAGGTCGTTCGCTCCAAG<br>CTGGGCTGTGTGCACGAACCCCCCGTTCAGCCCGACCGCTGCGCCTTATCCGGTAACTATCGTCTTGAGTCCAACCCGGT<br>AAGACACGACTTATCGCCACTGGCAGCAGCCACTGGTAACAGGATTAGCAGAGCGAGGTATGTAGGCGGTGCTACAGAGT<br>TCTTGAAGTGGTGGCCTAACTACGGCTACACTAGAAGGACAGTATTTGGTATCTGCGCTCTGCTGAAGCCAGTTACCTTC<br>GGAAAAAGAGTTGGTAGCTCTTGATCCGGCAAACAAACCACCGCTGGTAGCGGTGGTTTTTTTGTTTGCAAGCAGCAGAT<br>TACGCGCAGAAAAAAAGGATCTCAAGAAGATCCTTTGATCTTTTCTACGGGGTCTGACGCTCAGTGGAACGAAAACTCAC<br>GTTAAGGGATTTTGGTCATGAGATTATCAAAAAGGATCTTCACCTAGATCCTTTTAAATTAAAAATGAAGTTTTAAATCA<br>ATCTAAAGTATATATGAGTAAACTTGGTCTGACAGTTACCAATGCTTAATCAGTGAGGCACCTATCTCAGCGATCTGTCT<br>ATTTCGTTCATCCATAGTTGCCTGACTCCCCGTCGTGTAGATAACTACGATACGGGAGCGCTTACCATCTGGCCCCAGTG<br>CTGCAATGATACCGCGAGACCCACGCTCACCGGCTCCAGATTTATCAGCAATAAACCAGCCAGCCGGAAGGGCCGAGCGC<br>AGAAGTGGTCCTGCAACTTTATCCGCCTCCATTCAGTCTATTAATTGTTGCCGGGAAGCTAGAGTAAGTAGTTCGCCAGT<br>TAATAGTTTGCGCAACGTTGTTGGCATTGCTACAGGCATCGTGGTGTCACTCTCGTCGTTTGGTATGGCTTCATTCAGCT<br>CCGGTTCCCAACGATCAAGGCGAGTTACATGATCCCCCATGTTGTGCAAAAAAGCGGTTAGCTCCTTCGGTCCTCCGATC<br>GTTGTCAGAAGTAAGTTGGCCGCAGTGTTATCACTCATGGTTATGGCAGCACTGCATAATTCTCTTACTGTCATGCCATC<br>CGTAAGATGCTTTTCTGTGACTGGTGAGTACTCAACCAAGTCATTCTGAGAATAGTGTATGCGGCGACCGAGTTGCTCTT |

| | |
|---|---|
| | GCCCGGCGTCAATACGGGATAATAGTGTATCACATAGCAGAACTTTAAAAGTGCTCATCATTGGAAAACGTTCTTCGGGG<br>CGAAAACTCTCAAGGATCTTACCGCTGTTGAGATCCAGTTCGATGTAACCCACTCGTGCACCCAACTGATCTTCAGCATC<br>TTTTACTTTCACCAGCGTTTCTGGGTGAGCAAAAACAGGAAGGCAAAATGCCGCAAAAAAGGGAATAAGGGCGACACGGA<br>AATGTTGAATACTCATACTCTTCCTTTTTCAATGGGTAATAACTGATATAATTAAATTGAAGCTCTAATTTGTGAGTTTA<br>GTATACATGCATTTACTTATAATACAGTTTTTTAGTTTTGCTGGCCGCATCTTCTCAAATATGCTTCCCAGCCTGCTTTT<br>CTGTAACGTTCACCCTCTACCTTAGCATCCCTTCCCTTTGCAAATAGTCCTCTTCCAACAATAATAATGTCAGATCCTGT<br>AGAGACCACATCATCCACGGTTCTATACTGTTGACCCAATGCGTCTCCCTTGTCATCTAAACCCACACCGGGTGTCATAA<br>TCAACCAATCGTAACCTTCATCTCTTCCACCCATGTCTCTTTGAGCAATAAAGCCGATAACAAAATCTTTGTCGCTCTTC<br>GCAATGTCAACAGTACCCTTAGTATATTCTCCAGTAGATAGGGAGCCCTTGCATGACAATTCTGCTAACATCAAAAGGCC<br>TCTAGGTTCCTTTGTTACTTCTTCTGCCGCCTGCTTCAAACCGCTAACAATACCTGGGCCCACCACACCGTGTGCATTCG<br>TAATGTCTGCCCATTCTGCTATTCTGTATACACCCGCAGAGTACTGCAATTTGACTGTATTACCAATGTCAGCAAATTTT<br>CTGTCTTCGAAGAGTAAAAAATTGTACTTGGCGGATAATGCCTTTAGCGGCTTAACTGTGCCCTCCATGGAAAAATCAGT<br>CAAGATATCCACATGTGTTTTTAGTAAACAAATTTTGGGACCTAATGCTTCAACTAACTCCAGTAATTCCTTGGTGGTAC<br>GAACATCCAATGAAGCACACAAGTTTGTTTGCTTTTCGTGCATGATATTAAATAGCTTGGCAGCAACAGGACTAGGATGA<br>GTAGCAGCACGTTCCTTATATGTAGCTTTCGACATGATTTATCTTCGTTTCCTGCAGGTTTTTGTTCTGTGCAGTTGGGT<br>TAAGAATACTGGGCAATTTCATGTTTCTTCAACACTACATATGCGTATATATACCAATCTAAGTCTGTGCTCCTTCCTTC<br>GTTCTTCCTTCTGTTCGGAGATTACCGAATCAAAAAAATTTCAAAGAAACCGAAATCAAAAAAAAGAATAAAAAAAAAAT<br>GATGAATTGAATTGAAAAGCTAGCTTATCGATGATAAGCTGTCAAAGATGAGAATTAATTCCACGGACTATAGACTATAC<br>TAGATACTCCGTCTACTGTACGATACACTTCCGCTCAGGTCCTTGTCCTTTAACGAGGCCTTACCACTCTTTTGTTACTC<br>TATTGATCCAGCTCAGCAAAGGCAGTGTGATCTAAGATTCTATCTTCGCGATGTAGTAAAACTAGCTAGACCGAGAAAGA<br>GACTAGAAATGCAAAAGGCACTTCTACAATGGCTGCCATCATTATTATCCGATGTGACGCTGCAGCTTCTCAATGATATT<br>CGAATACGCTTTGAGGAGATACAGCCTAATATCCGACAAACTGTTTTACAGATTTACGATCGTACTTGTTACCCATCATT<br>GAATTTTGAACATCCGAACCTGGGAGTTTTCCCTGAAACAGATAGTATATTTGAACCTGTATAATAATATATAGTCTAGC<br>GCTTTACGGAAGACAATGTATGTATTTCGGTTCCTGGAGAAACTATTGCATCTATTGCATAGGTAATCTTGCACGTCGCA<br>TCCCCGGTTCATTTTCTGCGTTTCCATCTTGCACTTCAATAGCATATCTTTGTTAACGAAGCATCTGTGCTTCATTTTGT<br>AGAACAAAAATGCAACGCGAGAGCGCTAATTTTTCAAACAAAGAATCTGAGCTGCATTTTTACAGAACAGAAATGCAACG<br>CGAAAGCGCTATTTTACCAACGAAGAATCTGTGCTTCATTTTTGTAAAACAAAAATGCAACGCGACGAGAGCGCTAATTT<br>TTCAAACAAAGAATCTGAGCTGCATTTTTACAGAACAGAAATGCAACGCGAGAGCGCTATTTTACCAACAAAGAATCTAT<br>ACTTCTTTTTTGTTCTACAAAAATGCATCCCGAGAGCGCTATTTTTCTAACAAAGCATCTTAGATTACTTTTTTTCTCCT<br>TTGTGCGCTCTATAATGCAGTCTCTTGATAACTTTTTGCACTGTAGGTCCGTTAAGGTTAGAAGAAGGCTACTTTGGTGT<br>CTATTTTCTCTTCCATAAAAAAAGCCTGACTCCACTTCCCGCGTTTACTGATTACTAGCGAAGCTGCGGGTGCATTTTTT<br>CAAGATAAAGGCATCCCCGATTATATTCTATACCGATGTGGATTGCGCATACTTTGTGAACAGAAAGTGATAGCGTTGAT<br>GATTCTTCATTGGTCAGAAAATTATGAACGGTTTCTTCTATTTTGTCTCTATATACTACGTATAGGAAATGTTTACATTT<br>TCGTATTGTTTTCGATTCACTCTATGAATAGTTCTTACTACAATTTTTTTGTCTAAAGAGTAATACTAGAGATAAACATA<br>AAAAATGTAGAGGTCGAGTTTAGATGCAAGTTCAAGGAGCGAAAGGTGGATGGGTAGGTTATATAGGGATATAGCACAGA<br>GATATATAGCAAAGAGATACTTTTGAGCAATGTTTGTGGAAGCGGTATTCGCAATGGGAAGCTCCACCCCGGTTGATAAT<br>CAGAAAAGCCCCAAAAACAGGAAGATTGTATAAGCAAATATTTAAATTGTAAACGTTAATATTTTGTTAAAATTCGCGTT<br>AAATTTTTGTTAAATCAGCTCATTTTTTAACGAATAGCCCGAAATCGGCAAAATCCCTTATAAATCAAAAGAATAGACCG<br>AGATAGGGTTGAGTGTTGTTCCAGTTTCCAACAAGAGTCCACTATTAAAGAACGTGGACTCCAACGTCAAAGGGCGAAAA<br>AGGGTCTATCAGGGCGATGGCCCACTACGTGAACCATCACCCTAATCAAGTTTTTTGGGGTCGAGGTGCCGTAAAGCAGT<br>AAATCGGAAGGGTAAACGGATGCCCCCATTTAGAGCTTGACGGGGAAAGCCGGCGAACGTGGCGAGAAAGGAAGGGAAGA<br>AAGCGAAAGGAGCGGGGGCTAGGGCGGTGGGAAGTGTAGGGGTCACGCTGGGCGTAACCACCACACCCGCCGCGCTTAAT<br>GGGGCGCTACAGGGCGCGTGGGGATGATCCACTAGTACGGATTAGAAGCCGCCGAGCGGGTGACAGCCCTCCGAAGGAAG<br>ACTCTCCTCCGTGCGTCCTCGTCCTCACCGGTCGCGTTCCTGAAACGCAGATGTGCCTCGCGCCGCACTGCTCCGAACAA<br>TAAAGATTCTACAATACTAGCTTTTATGGTTATGAAGAGGAAAAATTGGCAGTAACCTGGCCCCACAAACCTTCAAATGA<br>ACGAATCAAATTAACAACCATAGGATGATAATGCGATTAGTTTTTTAGCCTTATTTCTGGGGTAATTAATCAGCGAAGCG<br>ATGATTTTTGATCTATTAACAGATATATAAATGCAAAAACTGCATAACCACTTTAACTAATACTTTCAACATTTTCGGTT<br>TGTATTACTTCTTATTCAAATGTAATAAAAGTATCAACAAAAAATTGTTAATATACCTCTATACTTTAACGTCAAGGAGA<br>AAAAACCCCGGATCGGACTACTAGCAGCTGTAATACGACTCACTATAGGGAATATTAAGCTTGGTACCctcgagATGGAT<br>GTATTCATGAAAGGACTTTCAAAGGCCAAGGAGGGAGTTGTGGCTGCTGCTGAGAAAACCAAACAGGGTGTGGCAGAAGC<br>AGCAGGAAAGACAAAAGAGGGTGTTCTCTATGTAGGCTCCAAAACCAAGGAGGGAGTGGTGCATGGTGTGGCAACAGTGG<br>CTGAGAAGACCAAAGAGCAAGTGACAAATGTTGGAGGAGCAGTGGTGACGGGTGTGACAGCAGTAGCCCAGAAGACAGTG<br>GAGGGAGCAGGGAGCATTGCAGCAGCCACTGGCTTTGTCAAAAAGGACCAGTTGGGCAAGAATGAAGAAGGAGCCCCACA<br>GGAAGGAATTCTGGAAGATATGCCTGTGGATCCTGACAATGAGGCTTATGAAATGCCTTCTGAGGAAGGGTATCAAGACT<br>ACGAACCTGAAGCCGGCGCCCAGATCCACCGGTCGCCAACCGCAGTGAGCAAGGGCGAGGAGCTGTTCACCGGGGTGGTG<br>CCCATCCTGGTCGAGCTGGACGGCGACGTAAACGGCCACAAGTTCAGCGTCCGCGGCGAGGGCGAGGGCGATGCCACCAA<br>CGGCAAGCTGACCCTGAAGTTCATCTGCACCACCGGCAAGCTGCCCGTGCCCTGGCCCACCCTCGTGACCACCTTCGGCT<br>ACGGCGTGGCCTGCTTCAGCCGCTACCCCGACCACATGAAGCAGCACGACTTCTTCAAGTCCGCCATGCCCGAAGGCTAC<br>GTCCAGGAGCGCACCATCTCTTTCAAGGACGACGGTACCTACAAGACCCGCGCCGAGGTGAAGTTCGAGGGCGACACCCT<br>GGTGAACCGCATCGAGCTGAAGGGCATCGACTTCAAGGAGGACGGCAACATCCTGGGGCACAAGCTGGAGTACAACTTCA<br>ACAGCCACAACGTCTATATCACGGCCGACAAGCAGAAGAACGGCATCAAGGCTAACTTCAAGATCCGCCACAACGTTGAG<br>GACGGCAGCGTGCAGCTCGCCGACCACTACCAGCAGAACACCCCCATCGGCGACGGCCCCGTGCTGCTGCCCGACAACCA<br>CTACCTGAGCCATCAGTCCGCCCTGAGCAAAGACCCCAACGAGAAGCGCGATCACATGGTCCTGCTGGAGTTCGTGACCG<br>CCGCCGGGATTACACATGGCATGGACGAGCTGTACAAGTAANNNNNNNNNNNNNNNNNNNNNNNNNNgagctcTCTAGAG<br>GGCCGCATCATGTAATTAGTTATGTCACGCTTACATTCACGCCCTCCCCCCACATCCGCTCTAACCGAAAAGGAAGGAGT<br>TAGACAACCTGAAGTCTAGGTCCCTATTTATTTTTTTATAGTTATGTTAGTATTAAGAACGTTATTTATATTTCAAATTT<br>TTCTTTTTTTTCTGTACAGACGCGTGTACGCATGTAACATTATACTGAAAACCTTGCTTGAGAAGGTTTTGGGACGCTCG<br>AAGGCTTTAATTTGCGGCC |
